# Protein Structure Refinement Guided by Atomic Packing Frustration Analysis

**DOI:** 10.1101/2020.07.19.211169

**Authors:** Mingchen Chen, Xun Chen, Shikai Jin, Wei Lu, Xingcheng Lin, Peter G. Wolynes

## Abstract

Recent advances in machine learning, bioinformatics and the understanding of the folding problem have enabled efficient predictions of protein structures with moderate accuracy, even for targets when there is little information from templates. All-atom molecular dynamics simulations provide a route to refine such predicted structures, but unguided atomistic simulations, even when lengthy in time, often fail to eliminate incorrect structural features that would allow the structure to become more energetically favorable owing to the necessity of making large scale motions and overcoming energy barriers for side chain repacking. In this study, we show that localizing packing frustration at atomic resolution by examining the statistics of the energetic changes that occur when the local environment of a site is changed allows one to identify the most likely locations of incorrect contacts. The global statistics of atomic resolution frustration in structures that have been predicted using various algorithms provide strong indicators of structural quality when tested over a database of 20 targets from previous CASP experiments. Residues that are more correctly located turn out to be more minimally frustrated than more poorly positioned sites. These observations provide a diagnosis of both global and local quality of predicted structures, and thus can be used as guidance in all-atom refinement simulations of the 20 targets. Refinement simulations guided by atomic packing frustration turn out to be quite efficient and significantly improve the quality of the structures.

## 2 Introduction

Both experiment and theory show that the energy landscapes of proteins are funneled towards their solved structures. ^1^ While the unfolded molten globule may contain many frustrated interactions, the frustration level decreases during folding. The overall stability of folded proteins typically does not come from just a few hyperstable residues but instead a consistency of interaction is manifest throughout the structure. Traps in the folding process, typically, have much native structure but only a few incorrectly located residues which often have locally frustrated interactions.

A twin challenge to understanding how proteins fold is the problem of practically predicting protein structures from sequence. Obtaining biomolecular structures with high resolution is helpful for understanding many biomolecular processes and is a prerequisite for many structure-based therapeutic designs. ^2,3^ For a long time, experimentally determining the three dimensional atomic coordinates of proteins through crystallography, NMR and more recently cryo-Electron Microscopy has been the only reliable approach for most practical purposes. The fast development of bioinformatics and simulations lately involving deep learning techniques are however now commonly providing predictions of protein structures at modest resolution.^4,5^ To refine the resulting moderate resolution predictions into structural models comparable in accuracy and resolution to those determined by experiment remains nontrivial. All-atom molecular dynamics simulations provide an approach for such refinement, but the rugged and sometimes still locally frustrated nature of the energy landscape leads to traps from which it is hard to escape when using most sampling methods with atomic forcefields. For this reason, until recently it was even unclear whether all-atom simulations could refine protein structures to better resolution. Fortunately, a set of long time-scale refinement simulations performed with the Anton machine have demonstrated that the refinement using appropriate restraints can improve the quality of structure predictions,^6^ but, at the same time, simulating the protein motion without employing such restraints allows the ensemble of structures to deviate far away from the proper native structure. This is not surprising given the often subtle balance between energy and entropy in folding. ^7^

These results nevertheless give hope that increasing the time scale of refinement simulations could result in significantly improved refinements. Feig and his coworkers, however, have shown that with current computational time limitations, such a direct approach gives only marginally better refinement even when the simulation time scale has been greatly increased.^8^ They showed however that extensive sampling on the rugged energy landscape could be combined with Markov state modeling to further improve protein structure predictions so as to outperform other methods tested during the CASP13.^9^ The expense of such calculations unfortunately hinders the broad application of this approach. The time scale difficulty arises from the slow barrier-crossings involving both large-scale deformations and local side chain variations. These large scale motions are slow partly because of the high density of the protein interior that also encourage strong coupling between side-chain motions to lead to glassy behavior. The difficulties of traversing these barriers can be partially overcome by forcing motions along slow principal coordinates of motion of the protein. A good approximation to these modes can be found from coarse-grained models. This strategy brings significant improvements in structural accuracy. To address the difficulties of making barrier-crossings, the idea of enhancing atomic sampling along slow principal motions which are calculated from coarse-grained models finally brings significant improvements. ^10^

Another approach, of course is again to introduce restraints into atomistic simulations as was proved to work in obtaining more native-like configurations in Anton simulations. Most of the restraints come from different types of structural information of the initial coordinates. These should be soft restraints that preserve the folding funnel but that do not introduce such strong topological constraints such that rearrangements are impeded. ^11,12^ Using these in combination consistently improves the accuracy of predicted structures, but the improvements again are moderate. It is noteworthy, however, that the general frustration, even computed at a coarse-grained level generally is reduced. ^11^

Neither of the strategies we have described specifically addresses a core problem in refining protein structures: what are the good features that must be preserved during refinement calculations versus what are the bad features that need to be adjusted to achieve greater accuracy ? Local frustration analysis provides a way to do this. To better diagnose both the global and local quality of structures in their all-atom representations, we employ a scheme to compute an appropriate all-atom frustration index for a protein structure. ^13^ This analysis allows us to evaluate frustration both locally on each residue and globally over the tertiary structure. To locate where the most frustrated interactions are found in a protein, one gathers statistics about the local energy changes that occur when the protein sequence and structure are perturbed locally, thus generating a set of decoys for which the total energy of the protein is then recomputed including the steric repulsions. The steric repulsive forces are sensitive to even local perturbations in the protein backbones, because the side chains need to be repacked to accommodate for the fluctuations in backbones, thus making steric repulsions an important contribution to diagnosing structural quality. These steric repulsions are resolved in correct structures but are not resolved typically in incorrect structures. The resulting frustration index by including the repulsive forces is called by us the “atomic packing frustration” to contrast it with a more forgiving index that allows for a greater deformation of the protein framework. At the same time, the repulsive clashes make the atomic resolution algorithm including the repulsion too sensitive for detecting functionally relevant frustration, given that the strong fluctuating clashes coming from steric overlaps are easily removed by slightly relaxing the backbone in the dynamic functioning of the protein. In a parallel study that explores the relationship between frustration and functional outcomes, we have used the vander waals idea that the energetic costs incurred by the repulsions should be excluded as actually contributing only to the entropy^14^ in order to monitor stability and specificity in a way that more realistically accounts for protein flexibility. The resulting atomic resolution frustration measures (that eliminate the harsh repulsions) correlate well with the physiological functions of biomolecular complexes like binding, allostery, catalytic sites and association with ligands. ^15^

In the present paper, we show that the atomic packing frustration which is indeed sensitive to steric clashes correlates well with the accuracy of the local placement of each residue within a protein, and the residues with good structural quality are usually more minimally frustrated from the point of the view of packing. The global frustration level of a protein structural model also correlates well with the global quality of the model, and models having more native-like poses usually have a larger number of minimally frustrated steric interactions. The capability of diagnosing both the local and global structural quality of modeled structures offers a way to select appropriate restraints in all-atom refinement simulations by constraining only the motions of the minimally frustrated residues found in the initial predicted model while allowing those regions with frustrated packing to readjust by roaming more widely. We will show that structures from refinement simulations using frustration guided restraints display significantly improved structural accuracy over the initial poses using simulations that correspond to only 50 ns of annealing. The poses after refinement were less frustrated than they are when they start out.

## 3 Methods

### 3.1 Definition of packing frustration

To capture the local stability of the placement of a single amino acid based on its interaction with neighboring residues, we introduce a new frustration index. Frustration indices are obtained by monitoring the energy differences between the protein in its native conformation and a set of “decoy” states. We collect statistics about these local energy contributions by perturbing both the protein sequence and local structure. To generate a set of decoys, the amino acid pairs which form the contacts in the crystal structures are virtually mutated to other amino acid pairs. The sequence spaces are randomly sampled according to the frequency distribution of the native amino acids each taking on different side chain conformations. Given the recomputed energy of 400 distributed decoys for each contact, we can thus construct a histogram of the energy of these decoys and compare them with the native energy, *E*_0_. The frustration index including an interacting pair of the amino acids *i, j* is defined as the Z score of the native pair’s energy compared with the variance of the pair’s contribution in the corresponding decoys:

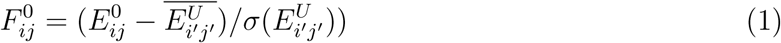

Here 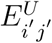 is the pairwise interaction energy of the decoy, 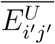 is the mean and 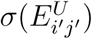 is the corresponding variance of energies. The frustration index is a site-specific measure of the energetic fitness compared with the set of all other possible decoys. The frustration index is thus the local energy gap between the native state and the decoys normalized the variance of its distribution. This measure is equivalent to that used in describing the folding funnel of evolved proteins which scales the global energy to the glass formation temperature. ^1^ If the frustration index of a pair is negative and significantly large in magnitude, the specific contact formed by that pair should be almost the most stable among all of the other alternatives and is unlikely to change during functional motions. If the frustration index is low, the alternative local structures become energetically accessible. The local frustration index evaluates the role of contacts in building the globally funneled folding energy landscape. As discussed in Ferreiro et al, ^15^ contacts can be classified as being minimally frustrated, highly frustrated or as being neutrally frustrated in accordance with their frustration level. In this study, contacts that have a positive frustration index above 0.5 are identified as being highly frustrated interactions, while contacts having a frustration index below -2.5 are identified as being minimally frustrated contacts.

We used the all-atom forcefield from Rosetta^16^ to compute localized packing frustration indices. To obtain the decoy ensemble for residue contacts, we randomized both the residue identities and locations of the residues in contact as was done previously when using the AWSEM coarse grained energy function. ^15^ We shuffled the protein sequence randomly and repacked these shuffled sequences onto the backbone whose coordinates stay unperturbed to make sure that only the side chains are re-packed. Then we performed a short Monte-Carlo relaxation to eliminate many of the most significant side chain clashes with the backbone fixed (see supporting information for the xml script). After these steps, all energies of the contacts are counted into the decoy ensemble 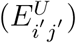. Sufficient statistics for each protein were collected after the decoy set was enriched by 300 shuffled sequences. Through omitting the shuffling step, we obtained the energies of native contacts in a similar fashion. Protein contacts are defined by the distances between *C*_*α*_ atoms of residues being below 10 Angstroms.

Based on the many-body construction of the fully atomic force field, the pairwise energy change between residue i and j, *E*_*ij*_ is determined by all the interactions that involve any of the two residues that are in contact. (Eq. 2).

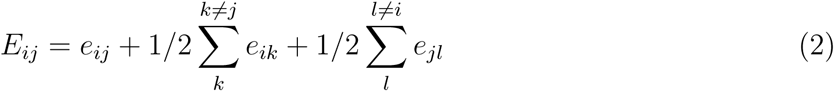

The direct interaction between residue i and j is defined as *e*_*ij*_. The auxiliary background interactions account for the many-body effects elicited by all the other residues of the protein except residue i or j. They are properly counted as half of 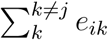 and 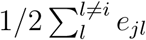. To compute the pairwise energy (*e*_*ij*_) in the relaxed pose, we have used the REF2015 version of the rosetta energy function^16^ along with a set of well-tested weights for each energy term.

Atomic packing frustration provides a way to estimate the localized steric frustration in proteins. To quantify the level of frustration for each specific residue, *F*_*i*_, we average all the pair frustration indices that involve the specific residue directly. The site averaged frustration *F*_*i*_ is used then to distinguish the minimally frustrated residues from those that can easily take on other geometries:

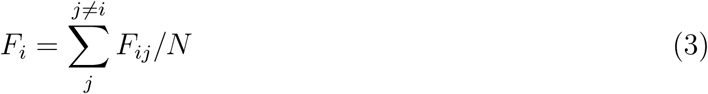

N is the total number of residues interacting with residue i.

### 3.2 Database of targets for tertiary structure refinement

The refinement targets studied in this paper came from the CASP12 refinement competition. The initial structures that had been previously predicted and the crystal structures for each target were downloaded from the official website of CASP12. When the website did not provide the crystal structures, we found them from the Protein Data Bank database ^17^ based on the PBD ID provided by the website. The submitted structures from the CASP12 competition were used as the refined structures in carrying out the evaluation. Here we used the submitted structures from the various groups to analyze the relationship between atomic packing frustration in models and global structural similarity to the correct structure.

### 3.3 Details of all-atom simulations censored by atomic frustration

The strategy is to refine a given initially predicted structure by reducing its localized packing frustration using all-atom explicit-solvent simulations. Here we used the CHARMM36m (Chemistry at Harvard Macromolecular Mechanics 36m) forcefield for the refinement simulations with a time step of 2.0 fs for 50ns. The total charge of the system was neutralized by adding Na and Cl ions to the solvent so that the final ionic concentration was 0.15 M. The simulations were performed in the constant particle number, volume, and temperature ensemble at 300K. The residual constraints were censored using the atomic packing frustration indices of this structure. The censored constraints were then applied as harmonic constraints on the positions of the initial C*α* atoms with a strength of 1000 kJ/mol.

### 3.4 Replica Exchange Molecular Dynamics (REMD)

To survey the all-atom free energy landscape, we performed parallel replica-exchange calculations.^18^ Replica exchange simulations were performed using the CHARMM36m forcefield over a range of temperatures at 250K, 270K, 290K, 310K, 330K, 350K, 370K, 390K, 410K, 430K, 450K and 470K. Exchange attempts were made every 0.2 ps, with acceptance rates ranging from 0.04 to 0.4. Each replica was equilibrated for 2 ns and then simulated for 30 ns for data collection.

### 3.5 Global Metrics of Structural and Side Chains Similarity

Three metrics were used to evaluate the structural similarity of refined structures to the corresponding crystal structures on the basis of the C*α* atoms. The C*α* RMSD value is computed using the root-mean-square deviation of the C*α* atoms of the refined structures from the positions of the corresponding C*α* atoms after alignment. The values of this metric were calculated using VMD. ^19^ Another metric we used was the GDT-TS score. It is defined as the average of the percentages of C*α* atoms falling within 1, 2, 4, and 8 angstrom of their correct locations upon alignment.^20^ The third metric to evaluate the structural similarity was Qw,^21^ defined by

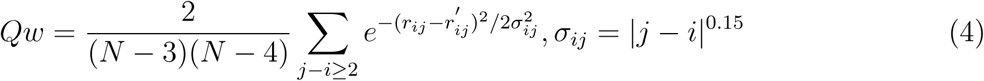

*r*_*ij*_ is the distance between the C*α* atom of residue index i and the C*α* atom of residue index j in the refined structure. 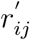 is the distance between the C*α* atoms of the corresponding residues in the native structures. *s*_*ij*_ is a sequence separation-dependent well width. The *Q*_*local*_ of residue i is defined as:

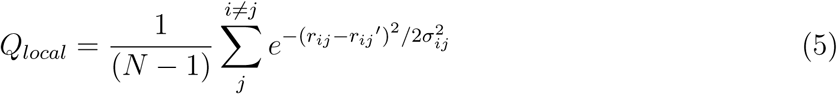

*r*_*ij*_, 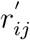, *s*_*ij*_ are defined as above.

The side chain similarity was defined as the fraction of the residues whose differences of both the *χ*1 and *χ*2 side chain angles in the refined structure and the corresponding crystal structure were less than 40 degrees.^11^

### 3.6 Blind selection based on logistic regression

To select structures with the best RMSD from all the refined structures without using the crystal structure information as input, we developed a logistic regression method similar to the one implemented in our previous paper.^10^ This logistic regression takes into account the energy terms from AWSEM^22^ and from Rosetta.^16,23–25^ AWSEM includes six terms:‘DSSP’, ‘P AP’, ‘Water’, ‘Burial’, ‘Helix’, ‘Electro’. Rosetta includes 16 terms: ‘fa atr’, ‘fa rep’, ‘fa sol’, ‘fa intra rep’, ‘fa intra sol xover4’, ‘lk ball wtd’, ‘fa elec’, ‘pro close’, ‘hbond sr bb’, ‘hbond lr bb’, ‘hbond bb sc’, ‘hbond sc’, ‘omega’, ‘fa dun’, ‘p aa pp’, ‘rama prepro’. These 22 energy terms are the feature inputs to the logistic regression. To have meaningful coefficients, all these quantitative features were normalized to have mean 0 and standard deviation 1 before the regression. The structures of TR884 were used as the training set for regression which classified the top 100 structures evaluated by RMSD out of the total 2500 structures as being good (y=1) and the rest as being bad (y=0). The logistic regression was trained to tune the weights(w) and intercept c that minimize the following cost function.

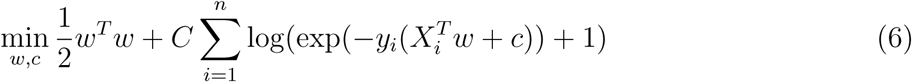

where C=1

## 4 Results

### 4.1 1: Correlation between global frustration and structural quality

The principle of minimal frustration suggests that a protein has evolved to minimize the energetic conflicts that need to be resolved when folding towards the native conformation. ^15^ Thus, the more native-like configurations generally possess more minimally frustrated interactions than the molten globule just as the final native configuration does. The frustration level of a given structure is determined by the number of minimally frustrated interactions it contains as well as by the number of highly frustrated interactions that it retains. To quantify the relationship between overall frustration level and global structural quality, here we calculated the atomic packing frustration for the submitted structures from the various groups for the 20 targets in the CASP12 refinement exercise.

The Qw value quantifies the global structural similarity between an input structure and the crystal structure. Fig.1 shows the structures submitted in the CASP12 refinement exercise along with crystal structures for four cases. The frustration patterns can discriminate the native structures from near native-like configurations (Fig.1, S1, S2, S3 and S4). When compared with the submitted structures for each target, one finds the native structure almost always has both the most minimally frustrated interactions and the fewest highly frustrated interactions. When approaching the native conformations, one finds the more native-like structures ,i.e. those, with a higher q value have again more minimally frustrated interactions and fewer highly frustrated interactions. The structures with extremely low structural quality (i.e. Q *<* 0.5, blue dots in Fig.1) have few minimally frustrated interactions. Sometimes structures with still reasonable similarity to the native configuration are not as minimally frustrated as other structures that have similar quality (Fig.S2A, S2C). In general, however the global statistics of atomic packing frustration over a protein structure does reflect the structural quality of a model quite well.

**Figure 1:**
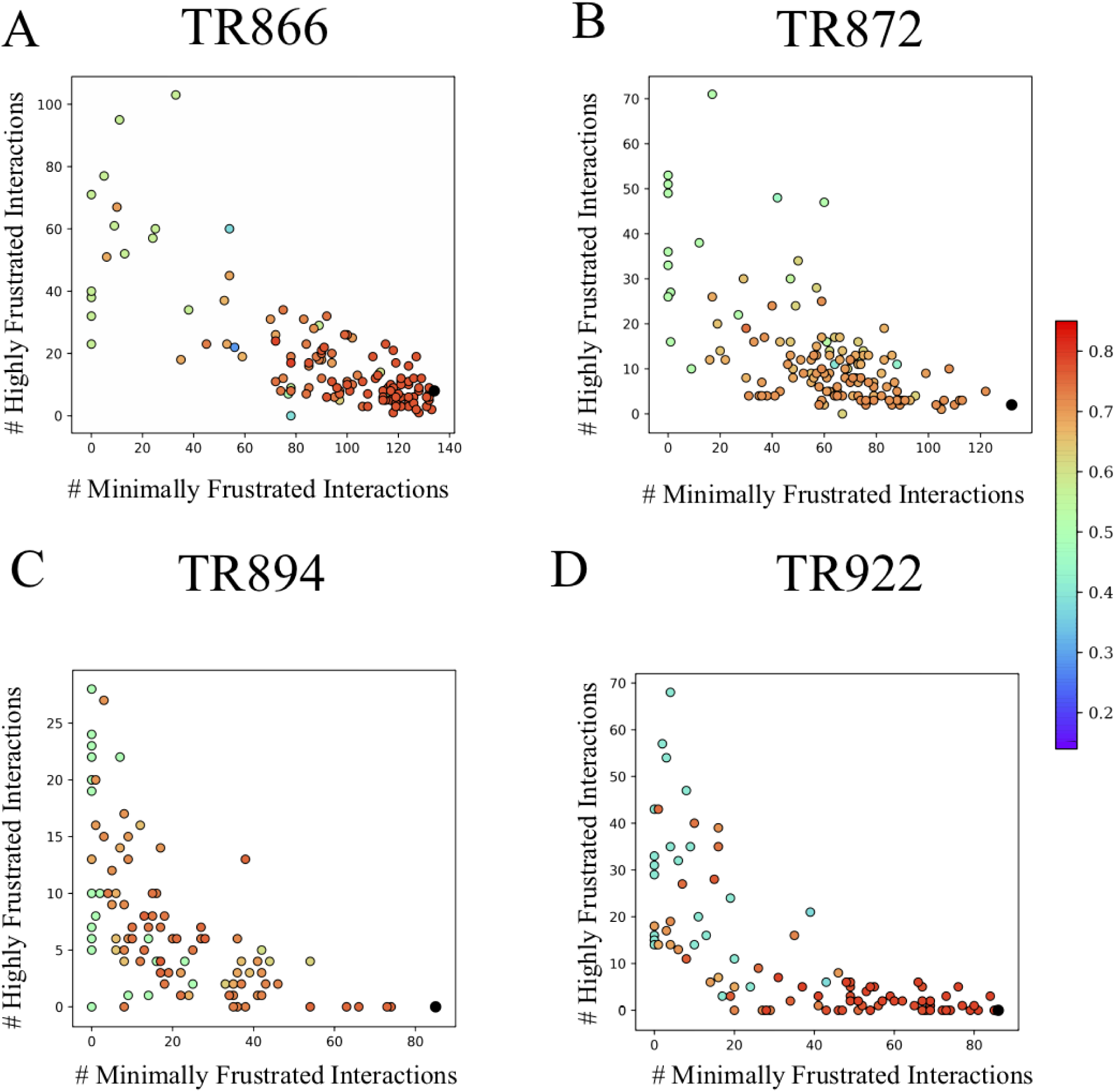
Results for four example targets show the correlation between frustration pattern and global similarity of the submitted refined structures and the corresponding crystal structures. A) TR866,B) TR872, C) TR894, D) TR922. All structures were from CASP12 groups. In each example, the frustration pattern was evaluated using the number of minimally frustrated interactions and the number of highly frustrated interactions. The global quality of each structure was evaluated using the Qw value between it and the crystal structure (shown in different colors corresponding to Qw), The points corresponding to crystal structures are colored in black.

### 4.2 Correlation between local structural quality and frustration pattern

During the process of protein folding, some native contacts readily form at the early stages, preceding the formation of the nucleus for the folding process, while some contacts in the native state require more time to form and do so only as the temperature is lowered. Those regions frequently pack with non-native interactions which are usually highly frustrated. To interpret the relationship between the residual frustration and the local structural quality, here we used a local measure of the quality of residue placement *Q*_*local*_ to quantify the local structural quality of the location of a residue in its tertiary environment. The residual frustration of each amino acid in sequence obtained from atomic packing frustration analyses can thus be compared with the correctness of the position of that residue as measured by *Q*_*local*_. Fig.2 shows the relationship between local frustration index and the local structural similarity of the initially predicted structure to the crystal structure for three targets in CASP12. The majority of the residues with move strongly negative frustration indices also display better structural quality locally. At the same time, these residues with lower local structural quality are highly frustrated in their local environment.

**Figure 2:**
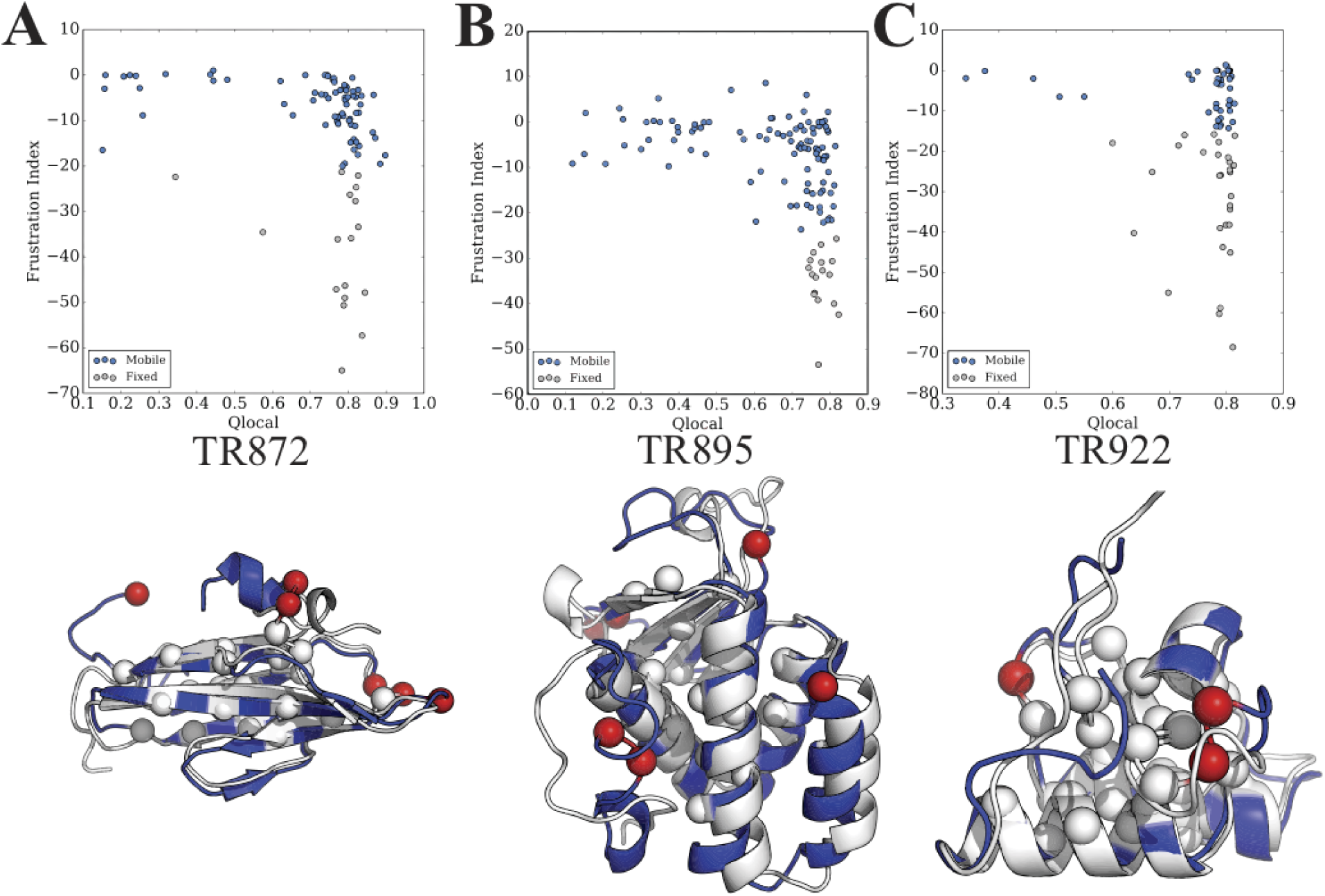
Three examples show the relationship between the frustration pattern and local structural accuracy of the initially predicted structure. A) TR872, B) TR895, C) TR922. The upper panel shows the relationship between the local frustration pattern evaluated by frustration indices and local structural similarity by *Q*_*local*_. In the lower panel, the initially predicted structure, colored by blue, was aligned to the crystal structure colored by white. The most minimally frustrated residues in the initially predicted structure which were constrained in refinement simulations are shown in white spheres. The most highly frustrated residues in the initially predicted structure are shown in red spheres.

In general, the local structural quality decreases with increasing local residue frustration as derived from packing frustration analysis of a structure. The residues with significantly large negative frustration indices form more native-like contacts with their surrounding residues. The residues which form non-native contacts or fewer contacts than the crystal structures have higher positive frustration indices. Some residues that display high frustration levels can still be quite native-like, but those residues are largely exposed on the surface. They are involved in function to locate binding sites, or key residues for allosteric effects. This is in harmony with the observation that protein surfaces are usually more enriched in frustrated interactions. ^15^ Based on the relationship between the frustration pattern and the structural quality locally, we have developed a refinement scheme that uses all-atom simulations that have constrained the motions of only the minimally frustrated residues to stay near their original positions found in the prediction. The bottom panel of Fig.2 shows both the minimally frustrated residues constrained in our test set of refinement simulations according to this protocol and that are for reference the most highly frustrated residues of the initial unrefined structures.

### 4.3 Overview of Atomic packing Frustration-Guided Refinement of CASP12 Targets

We summarize the results of our refinement protocol in Fig.3, Fig.S22, and Table.1. We compare the structural quality of the structures achieved using different refinement methods based on the RMSD as a quality metric (Fig.3). In Fig 3, the RMSD values of the refined structures from the‘atomic packing frustration-guided refinement’ are shown as red squares, whereas those of the initial models are shown as black squares. We also show the RMSD values of the refined structures from the Seok group^26^(blue squares), from the Feig Group^8^(yellow squares) and from the ‘PC-guided refinement’^10^(green squares).

**Table 1:**
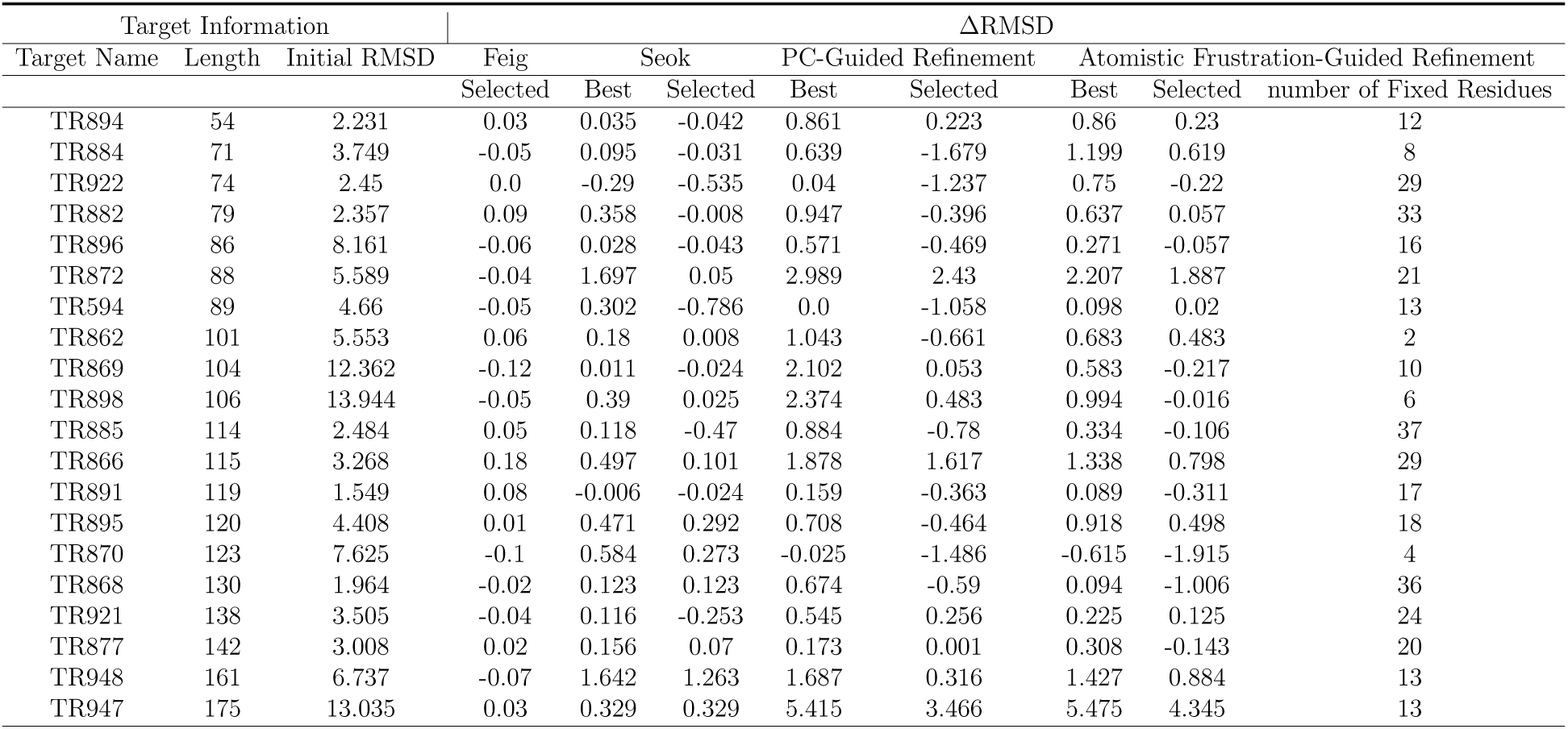
The Summary of Comparison between Different Refinement Methods

**Figure 3:**
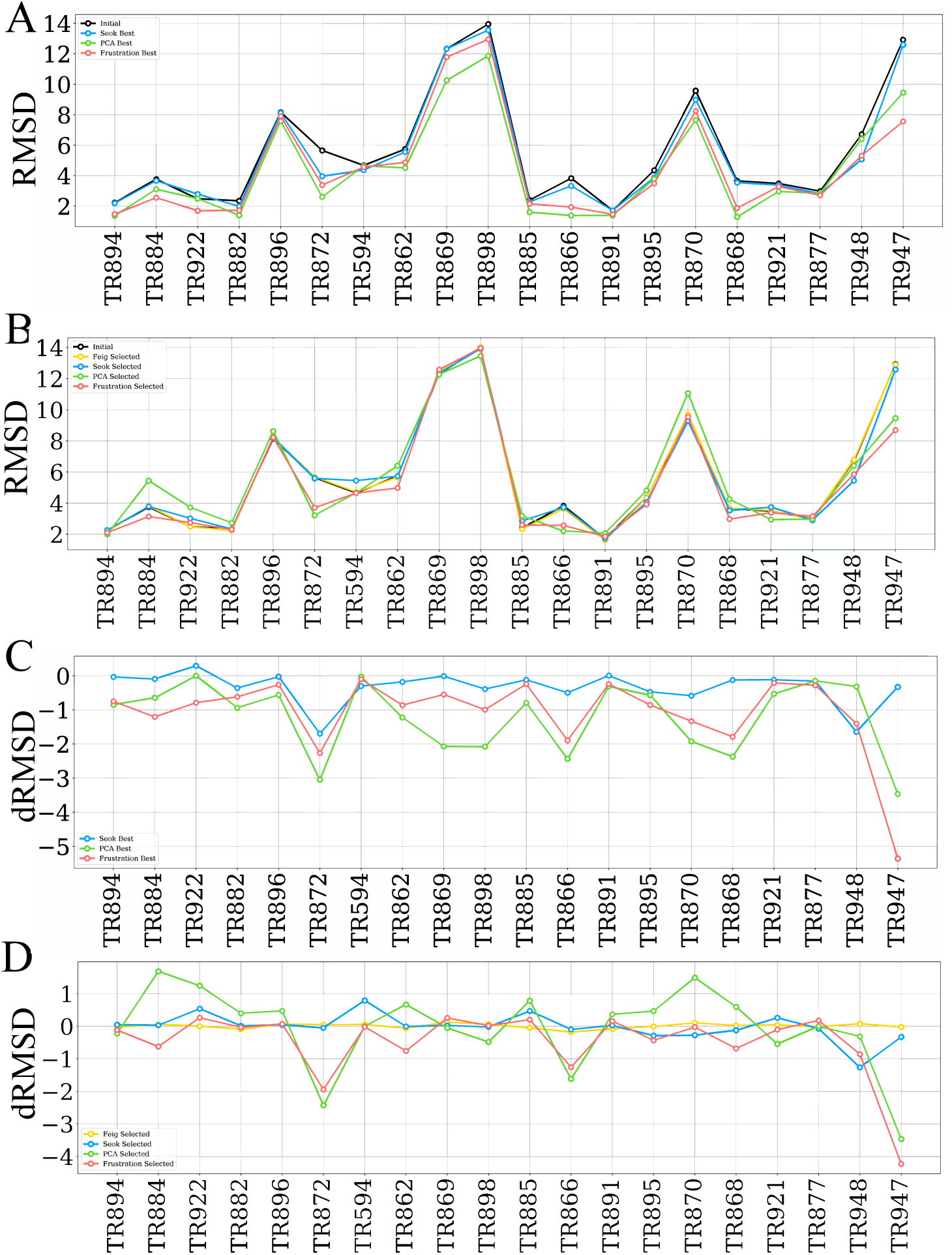
Summary of refinement results. A)the RMSD values of the initially predicted structures (”Initial”,Black) and the refined structures of the lowest RMSD values by different methods: Initial : Black, Seok Group: ^26^ Blue, PC-guided refinement: Green, Atomic Packing Frustration Guided Refinement: Red. B)the RMSD values of initially predicted structures and the selected structures from different refinement methods:Initial : Black, Feig Group: ^8^ Yellow, Seok Group: Blue, PC-guided refinement: Green, Atomic Packing Frustration Guided Refinement: Red.C)the ΔRMSD between initially predicted structures and the refined structures of lowest RMSD values by different methods:Seok Group: Blue, PC-guided refinement: Green, Atomic Packing Frustration Guided Refinement: Red. D)The Δrmsd between initially predicted structures and selected structures provided by different refinement schemes : Feig Group: Yellow, Seok Group: Blue, PC-guided refinement: Green, Atomic Packing Frustration Guided Refinement: Red. Feig group didn’t publish their refined structures of the lowest RMSD values.

The refined structures from the ‘atomic packing frustration-guided refinement’ have higher structural quality than the initially predicted models for each target. In contrast, the structural quality of the refined structures provided by the Seok group and the ‘PC-guided refinement’ is sometimes worse than the structural quality of the initially predicted model. This was seen for TR922. Though each refinement scheme improved the structural quality compared to the initially predicted models for TR898, TR895, TR877, and TR947, the ‘atomic packing frustration-guided refinement’ outperforms the other refinement methods.

Besides comparing the refined structures with the minimum RMSD values, we also compared the RMSD values of the refined structures that were obtained using different refinement schemes through blind selection in Fig.3C, Fig.3D, and Table.1. Based on the percentage of structures that were improved, the frustration informed refinement performs somewhat better than other refinement methods. The selected structures picked by our scheme were improved in structural quality relative to the initial models for 11 of 20 targets, whereas 10 of 20 targets were improved in the selected structures reported by Seok,^26^ 9 of 20 targets were improved as reported by Feig,^8^ and 9 of 20 targets were improved by ‘PC-guided refinement’.^10^ For those targets that were successfully improved to a lower RMSD value compared with the initially predicted models, the typical RMSD improvement was about 0.1 angstroms in the Seok and Feig studies. For the ‘atomic packing frustration-guided refinement’, the typical RMSD improvement is relatively larger than what was found using the Seok or the Feig methods, with the most significant improvement from the ‘atomic packing frustration-guided refinement’ being nearly 5 angstroms. Based on the average dRMSD values of the improved targets, our scheme yields a similar improvement as does‘PC-guided refinement’,^10^ which performs better than the earlier approach. ^8,26^

### 4.4 Atomic Packing Frustration-Guided Refinement Drives the Structure Towards it Native State

For each target, the ‘atomic packing frustration-guided refinement’ drives the protein closer towards its experimentally determined native state. To analyze the correlations between the change of the frustration pattern and the quality of the configurations, we compared the frustration patterns of the crystal structure to those of the initially predicted structures and the structures selected through logistic regression of the energetic components after collecting all the frustration indices for the component interactions.

For TR872 (Fig.4A), in the initially predicted model (middle panel), some contacts between the helix and its surrounding residues were highly frustrated. In contrast, these contacts become minimally frustrated in the refined structure (right panel) and are minimally frustrated in the crystal structure (left panel). For TR895 (Fig.4B), in the initially predicted structure (middle panel), the contacts formed between the loops are highly frustrated. In contrast, these contacts become neutrally frustrated in the resulting refined structure (right panel) and are in fact minimally frustrated in the crystal structures (left panel). For TR922 (Fig.4C), in the initially predicted model (middle panel), the contacts between the C terminal and its surrounding loop are highly frustrated, while these contacts become less highly frustrated in the structure after refinement (right panel), and the contacts are minimally frustrated in the correct crystal structures (left panel).

**Figure 4:**
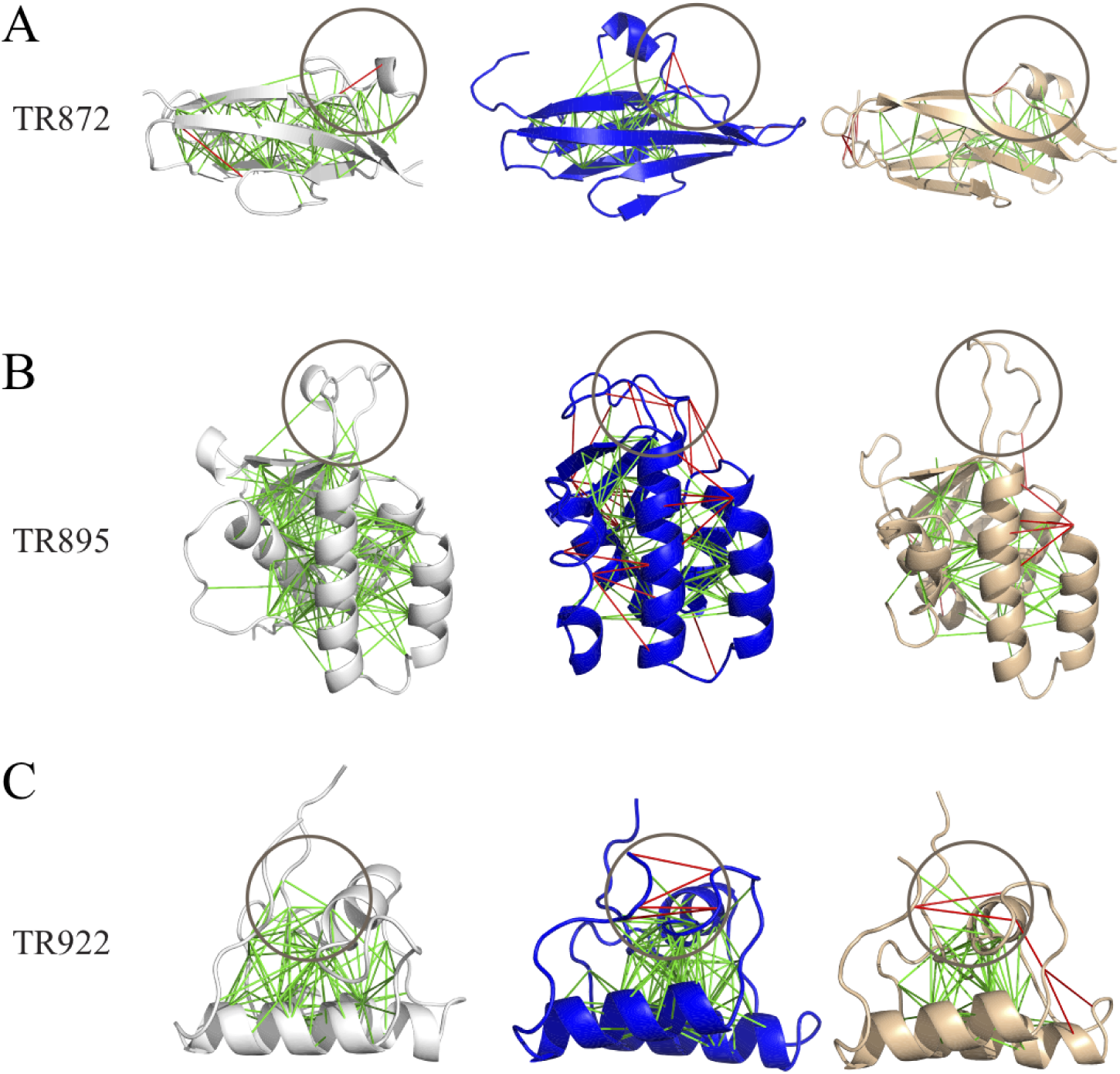
Differences of the frustration patterns between the crystal structures, the initially predicted structures and the refined structure obtained by blind selection. A) TR872,B) TR895, C) TR922. The left panel shows the frustration pattern of the crystal structures, the middle panel shows the frustration pattern of the initially predicted structures, and the right panel shows the frustration pattern of the selected structures from refinement. The green lines indicate minimally frustrated interactions and the red lines indicate highly frustrated interactions. The gray circle highlights the regions where the RMSD values and the amount of highly frustrated interactions decrease from those of the initially predicted structures, which suggests the effectiveness of atomic packing frustration guided refinement.

Similar trends were observed for most targets, except for those that showed minimal improvement in structural accuracy upon refinement(i.e. TR869, TR882, TR896, TR898 and TR891). For the refinement targets TR869, TR896 and TR898, the initially predicted structures deviate quite far from their corresponding crystal structures. Thus the derived constraints from frustration analyses of these initial structures still functioned poorly in guiding the structures through all-atom simulations. To understand this, we carried out allatom replica exchange molecular dynamics (REMD) simulations of TR872 to survey its free energy landscape. In the free energy surface of TR872 at 300K, the native-like basin (q ≥ 0.8) possesses more minimally frustrated interactions, but another basin far from its native state was observed at a Q around 0.55 that had fewer minimally frustrated interactions (Fig.S25). It is possible that this protein exhibits allostery and has multiple conformations in vivo. This set of results, however, clearly demonstrates the power of atomistic simulations censored by frustration analysis to improve input structures. By constraining the dynamics of the minimally frustrated residues in the input structure, atomic simulations are able adequately to sample over the motions of the less-minimally frustrated regions so as to minimize the total frustration level. As a result, the refined structures are usually more minimally frustrated than the initial forms and apparently are closer to the ground truth.

### 4.5 Improvement in Side chain Accuracy in Atomic Packing Frustration Guided Refinement

The RMSD value, Qw value, and GDT-TS score only reflect the accuracy of the backbones. Though the backbones restrict the orientations of side chains, rather slight differences arising from the backbone can lead to significant changes in the rotamers. Having these correct orientations of side-chains is generally considered important for describing binding with drugs, DNA, and catalysts, which are essential in various physiological functions.

Fig.5 shows the side chain accuracies for the *χ*1 and *χ*2 angles for all residues. The ‘atomic packing frustration-guided refinement’ greatly improves the accuracy of orientations of the side-chains in comparison with the input structure for all targets. This suggests that during the refinement process, the ‘atomic packing frustration-guided refinement’ not only improves the structural quality of the backbone, but also facilitates the correct native packing of side chains at the same time. This is consistent with the definition of the atomic packing frustration. The native packing of the side-chains eliminates steric clashes in comparsion with what is possible where the packing in highly non-native.

**Figure 5:**
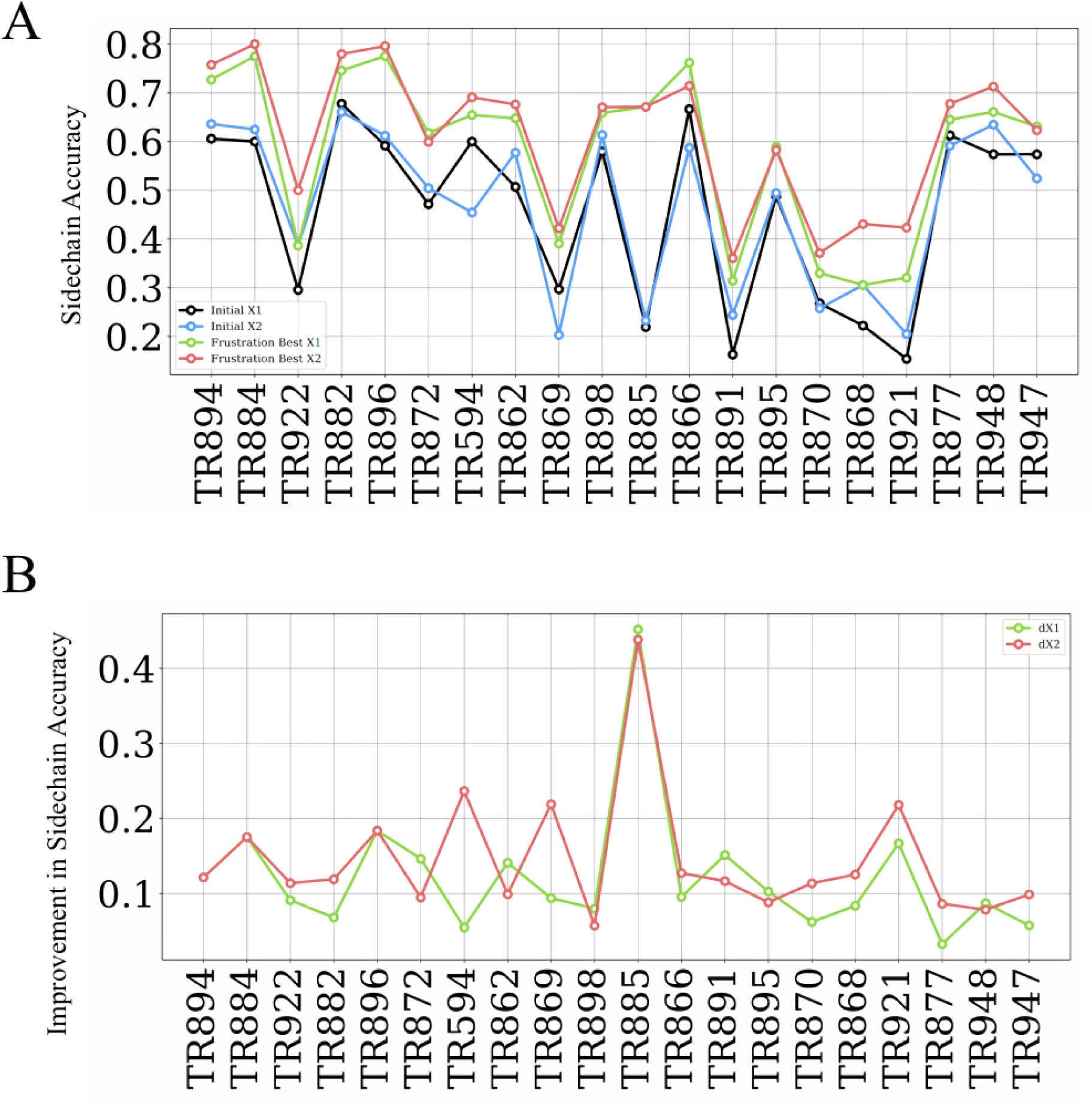
A summary of the X1 accuracy and X2 accuracy for all the residues. A)the X1 and X2 Accuracy of initially predicted structures and best structures from atomic packing frustration guided refinement,Black: X1 accuracy of the initially predicted structures, Blue: X2 accuracy of the initially predicted structures, Green: X1 accuracy of the best structures from atomic packing frustration guided refinement, Red: X2 accuracy of the best structures from atomic packing frustration guided refinement B)the difference in sidechain orientations accuracy between the best structures from atomic packing frustration guided refinement and initial structures.Green: Δ X1 accuracy, Red: Δ X2 accuracy.

**Figure 6:**
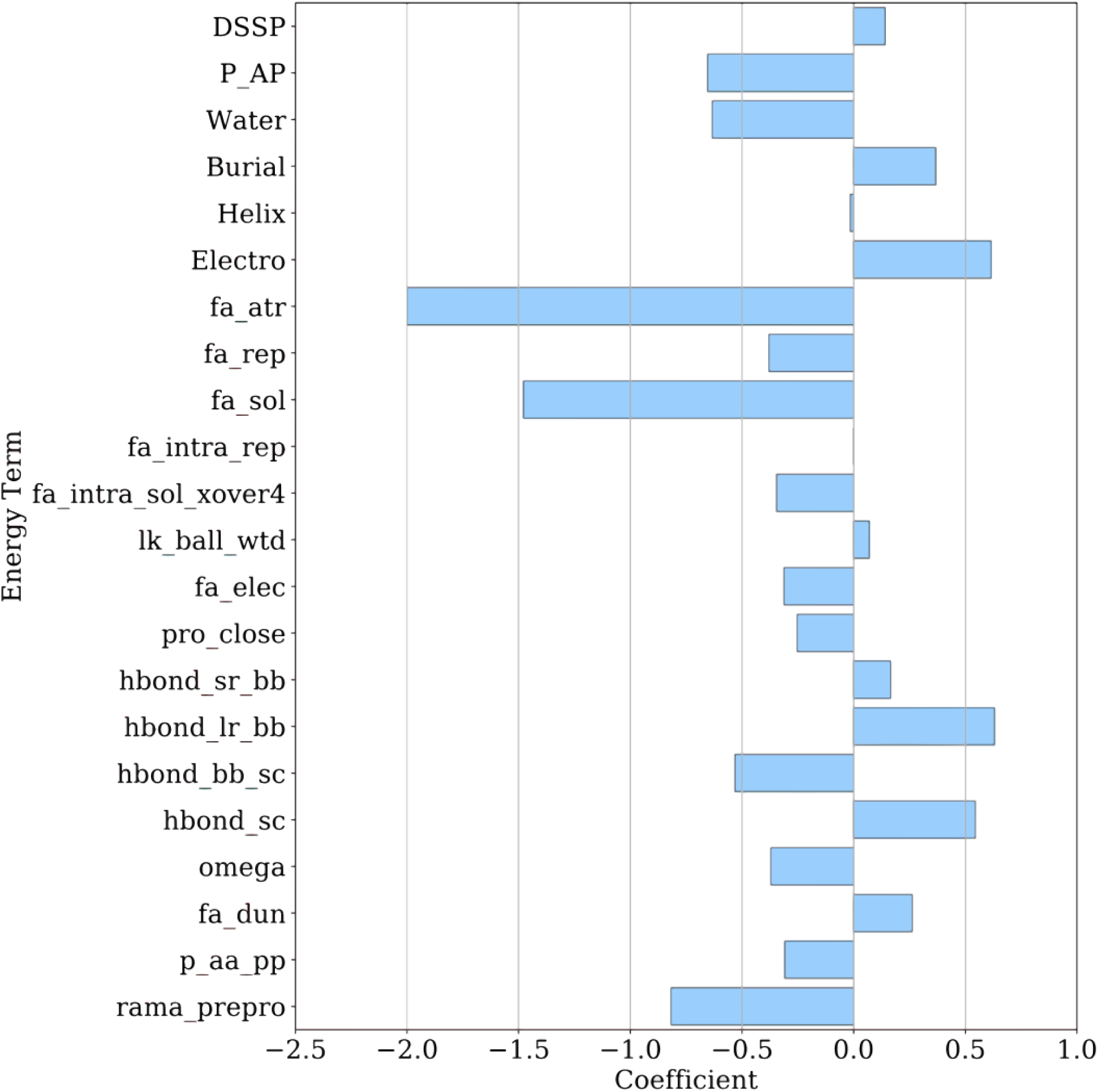
Coefficients of different features from AWSEM and Rosetta in linear regression selection

## 5 Discussion

### 5.1 The Roles of Different Energy Terms in Evaluating Structures

The top-scoring models obtained from atomic packing frustration guided refinement are better than those provided by other methods (in Fig.3C, Fig.3D and Table.1). The scheme used in the ‘atomic packing frustration-guided refinement’ unravels the roles of different parts of forcefield in protein folding and structure formation. The selection scheme uses 22 energy terms as input features to a logistic regression. These include all the energetic terms from AWSEM and from Rosetta. The absolute coefficients of these features reflect their contribution to accurate structure prediction when these features were normalized with mean 0 and standard deviation 1.

In AWSEM, the water term encodes the preference for a residue to form water-mediated or protein-mediated interactions, and the burial term indicates the inclination of one residue to become buried inside the protein or to stay on the surface.^22^ The P-AP term stabilizes the anti-parallel *β* conformation cooperatively, ^22^ and the electrostatic term stabilizes the folded state and destabilizes the unfolded states in the AWSEM system. ^27^ The high absolute coefficients of these features suggest their important roles in structure discrimination.

In terms of the energetic features from Rosetta, the attractive energy(fa atr), the solvation energy(fa sol), and the hydrogen-bonded energy(hbond lr bb, hbond sr bb, hbond bb sc) have relatively high absolute coefficients. This again suggests that these energy terms function differently in the final native state than they do in near native decoys.^16,23–25^

In this paper, we have illustrated how atomic packing frustration guided refinement drives the structure towards its most correctly configured state by the enhancement of its minimal frustration pattern. The current protocol improves structural quality efficiently by constraining only the minimally frustrated residues to be stably located in their predicted positions.

## 6 Acknowledgement

This work was initially supported by Grant R01 GM44557 from the National Institute of General Medical Sciences. Additional support was also provided by the D.R. Bullard-Welch Chair at Rice University, Grant C-0016. We thank the Data Analysis and Visualization Cyberinfrastructure funded by National Science Foundation Grant OCI-0959097, as well the Center for Theoretical Biological Physics sponsored by NSF Grant PHY-1427654.

## References

(1) Bryngelson, J. D.; Onuchic, J. N.; Socci, N. D.; Wolynes, P. G. Funnels, pathways, and the energy landscape of protein folding: a synthesis. Proteins: Structure, Function, and Bioinformatics 1995, 21, 167–195.

(2) Velanker, S. S.; Ray, S. S.; Gokhale, R. S.; Suma, S.; Balaram, H.; Balaram, P.; Murthy, M. Triosephosphate isomerase from Plasmodium falciparum:. the crystal struc-ture provides insights into antimalarial drug design. Structure 1997, 5, 751–761.

(3) Saphire, E. O.; Parren, P. W.; Pantophlet, R.; Zwick, M. B.; Morris, G. M.; Rudd, P. M.; Dwek, R. A.; Stanfield, R. L.; Burton, D. R.; Wilson, I. A. Crystal structure of a neutralizing human IGG against HIV-1: a template for vaccine design. science 2001, 293, 1155–1159.

(4) AlQuraishi, M. AlphaFold at CASP13. Bioinformatics 2019, 35, 4862–4865.

(5) Xu, J.; Wang, S. Analysis of distance-based protein structure prediction by deep learning in CASP13. Proteins: Structure, Function, and Bioinformatics 2019, 87, 1069–1081.

(6) Raval, A.; Piana, S.; Eastwood, M. P.; Dror, R. O.; Shaw, D. E. Refinement of protein structure homology models via long, all-atom molecular dynamics simulations. Proteins: Structure, Function, and Bioinformatics 2012, 80, 2071–2079.

(7) Shi, J.; Nobrega, R. P.; Schwantes, C.; Kathuria, S. V.; Bilsel, O.; Matthews, C. R.; Lane, T.; Pande, V. S. Atomistic structural ensemble refinement reveals non-native structure stabilizes a sub-millisecond folding intermediate of CheY. Scientific reports 2017, 7, 44116.

(8) Heo, L.; Feig, M. What makes it difficult to refine protein models further via molecular dynamics simulations? Proteins: Structure, Function, and Bioinformatics 2018, 86, 177–188.

(9) Heo, L.; Feig, M. High-accuracy protein structures by combining machine-learning with physics-based refinement. Proteins: Structure, Function, and Bioinformatics 2019,

(10) Lin, X.; Schafer, N. P.; Lu, W.; Jin, S.; Chen, X.; Chen, M.; Onuchic, J. N.; Wolynes, P. G. Forging tools for refining predicted protein structures. Proceedings of the National Academy of Sciences 2019, 116, 9400–9409.

(11) Chen, M.; Lin, X.; Lu, W.; Schafer, N. P.; Onuchic, J. N.; Wolynes, P. G. Template-guided protein structure prediction and refinement using optimized folding landscape force fields. Journal of chemical theory and computation 2018, 14, 6102–6116.

(12) Jin, S.; Chen, M.; Chen, X.; Bueno, C.; Lu, W.; Schafer, N. P.; Lin, X.; Onuchic, J. N.; Wolynes, P. G. Protein Structure Prediction in CASP13 using AWSEM-Suite. Journal of Chemical Theory and Computation 2020,

(13) Chen, J.; Schafer, N. P.; Wolynes, P. G.; Clementi, C. Localizing Frustration in Proteins Using All-Atom Energy Functions. The Journal of Physical Chemistry B 2019, 123, 4497–4504.

(14) Chandler, D.; Weeks, J. D.; Andersen, H. C. Van der Waals picture of liquids, solids, and phase transformations. Science 1983, 220, 787–794.

(15) Ferreiro, D. U.; Hegler, J. A.; Komives, E. A.; Wolynes, P. G. Localizing frustration in native proteins and protein assemblies. Proceedings of the National Academy of Sciences 2007, 104, 19819–19824.

(16) Alford, R. F.; Leaver-Fay, A.; Jeliazkov, J. R.; O’Meara, M. J.; DiMaio, F. P.; Park, H.; Shapovalov, M. V.; Renfrew, P. D.; Mulligan, V. K.; Kappel, K.; Others, The Rosetta all-atom energy function for macromolecular modeling and design. Journal of chemical theory and computation 2017, 13, 3031–3048.

(17) Berman, H. M.; Bourne, P. E.; Westbrook, J.; Zardecki, C. Protein Structure; CRC Press, 2003; pp 394–410.

(18) Hansmann, U. H. Parallel tempering algorithm for conformational studies of biological molecules. Chemical Physics Letters 1997, 281, 140–150.

(19) Humphrey, W.; Dalke, A.; Schulten, K.; Others, VMD visual molecular dynamics. Journal of molecular graphics 1996, 14, 33–38.

(20) Zhang, Y. Progress and challenges in protein structure prediction. Current opinion in structural biology 2008, 18, 342–348.

(21) Cho, S. S.; Levy, Y.; Wolynes, P. G. P versus Q: Structural reaction coordinates capture protein folding on smooth landscapes. Proceedings of the National Academy of Sciences 2006, 103, 586–591.

(22) Davtyan, A.; Schafer, N. P.; Zheng, W.; Clementi, C.; Wolynes, P. G.; Papoian, G. A. AWSEM-MD: protein structure prediction using coarse-grained physical potentials and bioinformatically based local structure biasing. The Journal of Physical Chemistry B 2012, 116, 8494–8503.

(23) Park, H.; Bradley, P.; Greisen Jr, P.; Liu, Y.; Mulligan, V. K.; Kim, D. E.; Baker, D.; DiMaio, F. Simultaneous optimization of biomolecular energy functions on features from small molecules and macromolecules. Journal of chemical theory and computation 2016, 12, 6201–6212.

(24) Leaver-Fay, A.; O’Meara, M. J.; Tyka, M.; Jacak, R.; Song, Y.; Kellogg, E. H.; Thompson, J.; Davis, I. W.; Pache, R. A.; Lyskov, S.; Others, Methods in enzymology; Elsevier, 013; Vol. 523; pp 109–143.

(25) O’Meara, M. J.; Leaver-Fay, A.; Tyka, M. D.; Stein, A.; Houlihan, K.; DiMaio, F.; Bradley, P.; Kortemme, T.; Baker, D.; Snoeyink, J.; Others, Combined covalent-electrostatic model of hydrogen bonding improves structure prediction with Rosetta. Journal of chemical theory and computation 2015, 11, 609–622.

(26) Lee, G. R.; Heo, L.; Seok, C. Simultaneous refinement of inaccurate local regions and overall structure in the CASP12 protein model refinement experiment. Proteins: Structure, Function, and Bioinformatics 2018, 86, 168–176.

(27) Tsai, M.-Y.; Zheng, W.; Balamurugan, D.; Schafer, N. P.; Kim, B. L.; Cheung, M. S.; Wolynes, P. G. Electrostatics, structure prediction, and the energy landscapes for protein folding and binding. Protein Science 2016, 25, 255–269.

