## Supplementary Information for "Protein Structure Refinement Guided by Atomic Packing Frustration Analysis"

### Supplementary Materials for Protein Structure Refinement Guided by Atomic Packing Frustration Analysis

### Contents

**S1 The evaluation of structural quality based on frustration pattern** **2**

**S2 Correlation between local structural quality and frustration pattern.** **6**

**S3 The change of sidechain accuracy during simulation** **12**

**S4 Refinements guided by atomic packing frustration guides the structures towards its experimentally determined native states** **17**

**S5 Comparing performance of atomic packing frustration guided refinement with other refinement methods by GDT-TS score** **23**

**S6 The free-energy landscapes of TR872 at temperature 300K** **27**

### S1 The evaluation of structural quality based on frustration pattern

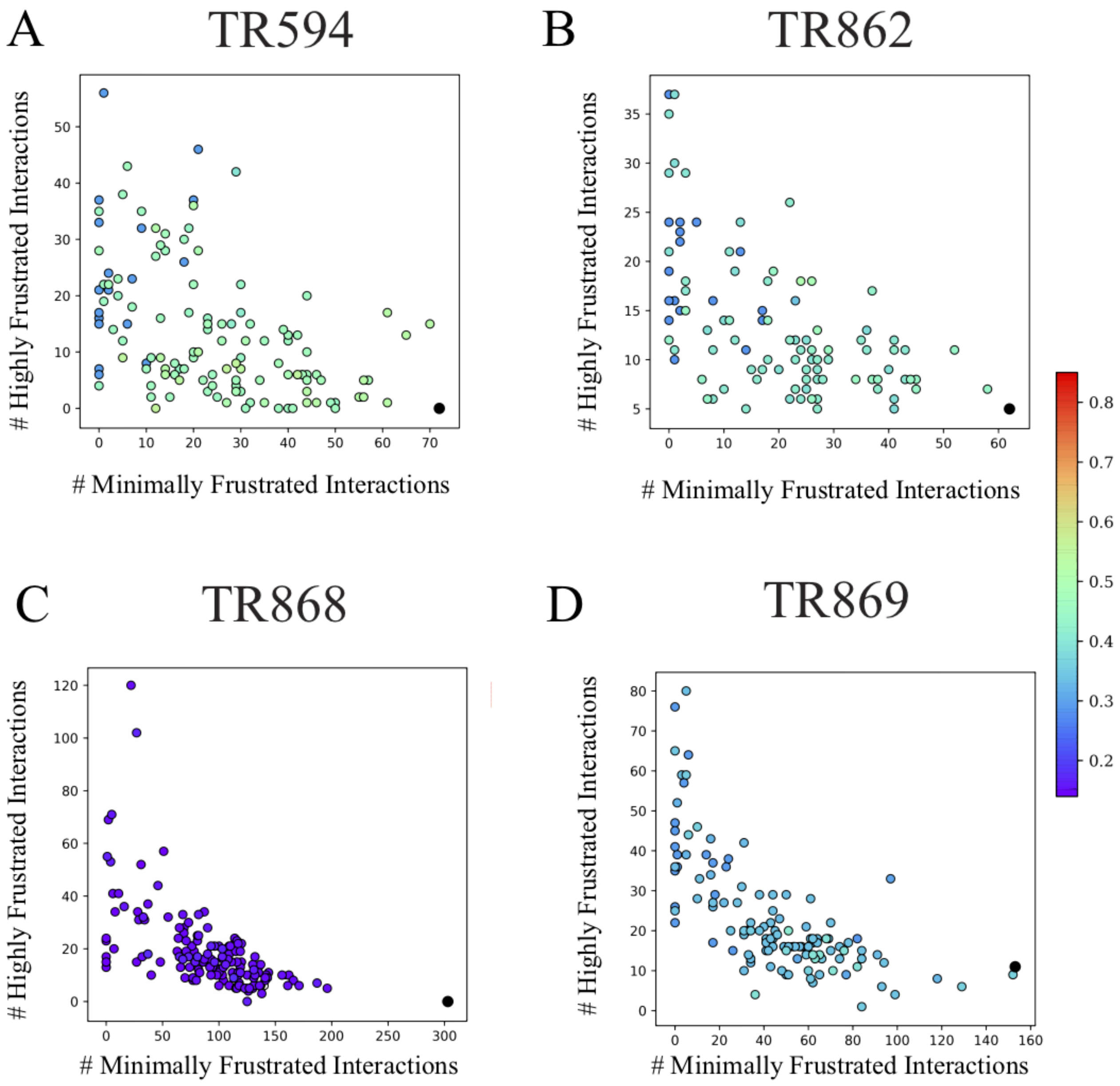


Figure S1: **Results for four example targets which show the correlation between frustration pattern and global similarity between the submitted refined structures and the corresponding crystal structures** A) TR594,B) TR862, C) TR868, D) TR869. All structures were from CASP12 groups. In each example, the frustration pattern was evaluated using the number of minimally frustrated interactions and the number of highly frustrated interactions. The global quality of each structure was evaluated using the Qw value between it and the crystal structure (shown in different colors corresponding to Qw), The points corresponding to crystal structures are colored in black.

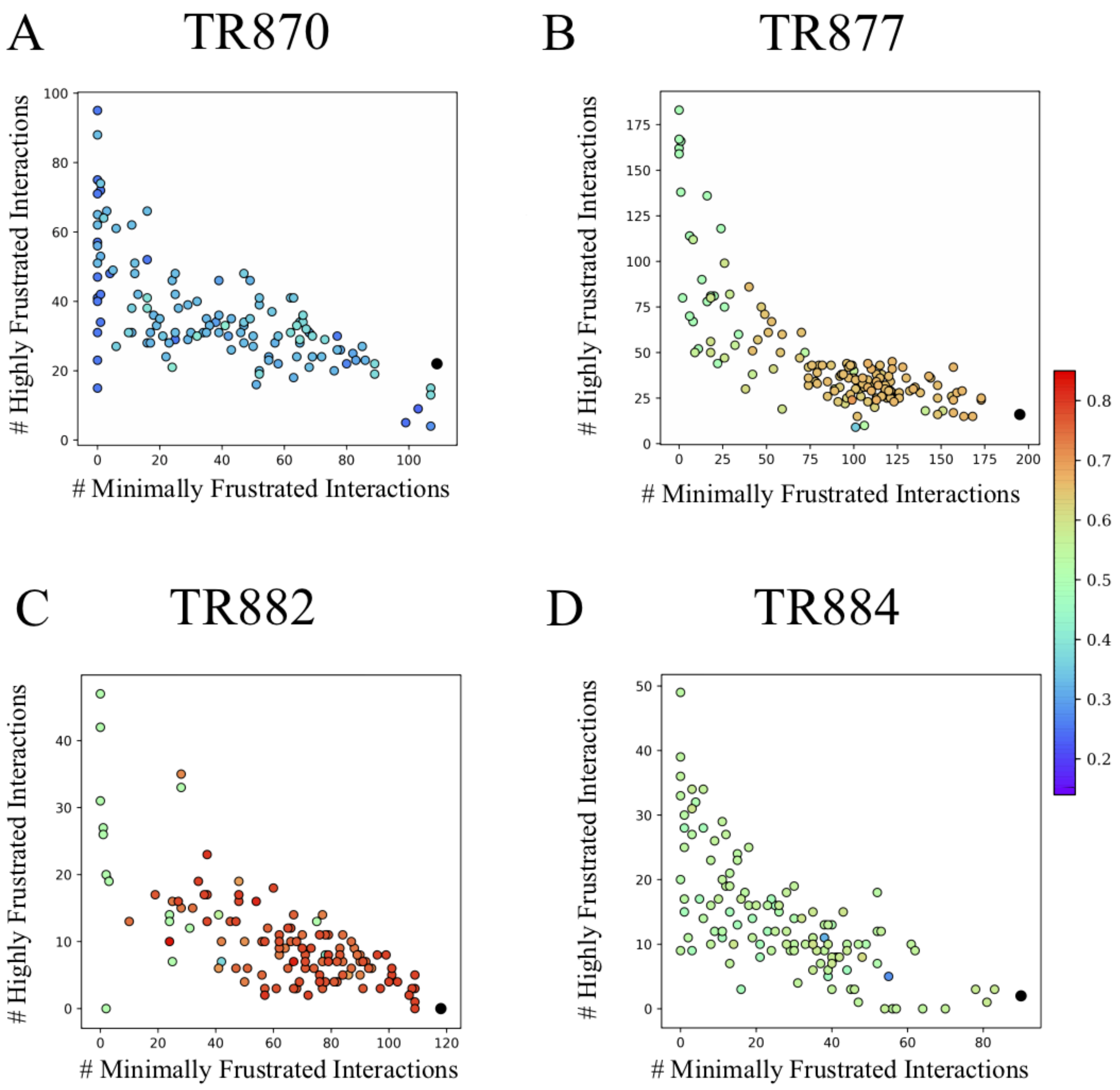


Figure S2: **Results for four example targets which show the correlation between frustration pattern and global similarity between the submitted refined structures and the corresponding crystal structures** A) TR870, B) TR877, C) TR882, D) TR884. All structures were from CASP12 groups. In each example, the frustration pattern was evaluated using the number of minimally frustrated interactions and the number of highly frustrated interactions. The global quality of each structure was evaluated using the Qw value between it and the crystal structure (shown in different colors corresponding to Qw), The points corresponding to crystal structures are colored in black.

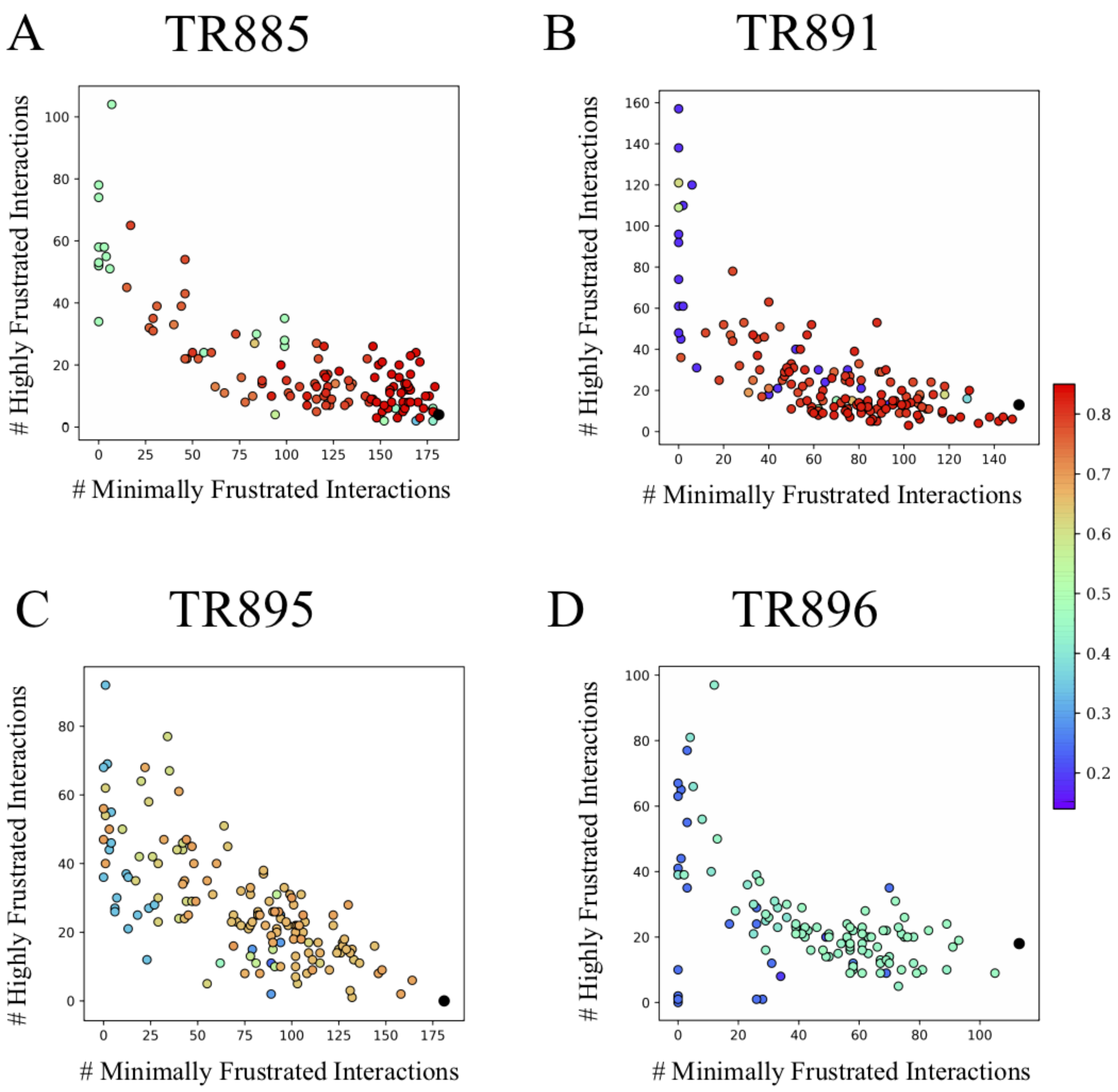


Figure S3: **Results for our example targets which show the correlation between frustration pattern and global similarity between the submitted refined structures and the corresponding crystal structures** A) TR885, B) TR891, C) TR895, D) TR896. All structures were from CASP12 groups. In each example, the frustration pattern was evaluated using the number of minimally frustrated interactions and the number of highly frustrated interactions. The global quality of each structure was evaluated using the Qw value between it and the crystal structure (shown in different colors corresponding to Qw), The points corresponding to crystal structures are colored in black.

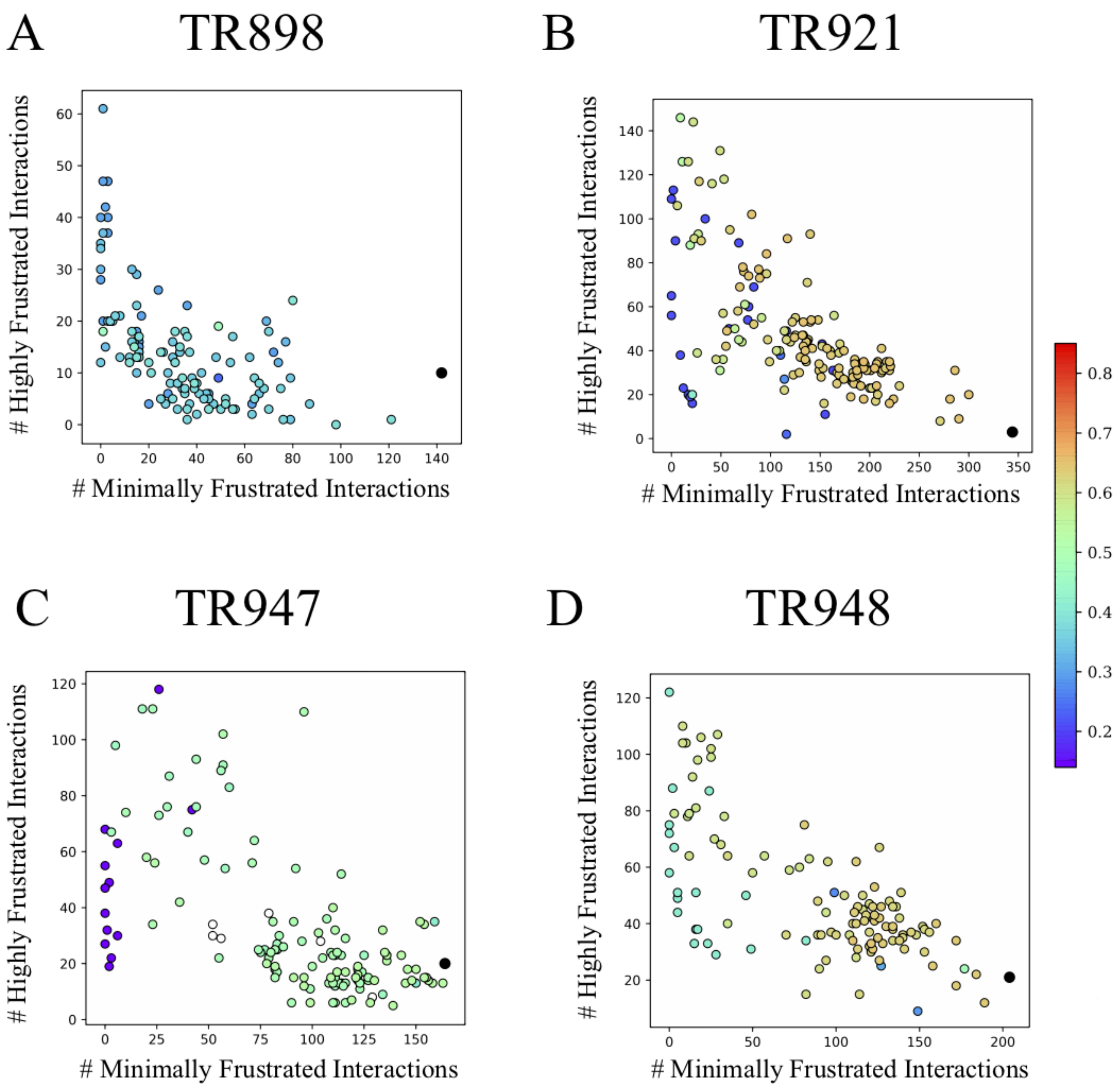


Figure S4: **Results for four example targets which show the correlation between frustration pattern and global similarity between the submitted refined structures and the corresponding crystal structures** A) TR898, B) TR921, C) TR947, D) TR948. All structures were from CASP12 groups. In each example, the frustration pattern was evaluated using the number of minimally frustrated interactions and the number of highly frustrated interactions. The global quality of each structure was evaluated using the Qw value between it and the crystal structure (shown in different colors corresponding to Qw), The points corresponding to crystal structures are colored in black.

### S2 Correlation between local structural quality and frustration pattern.

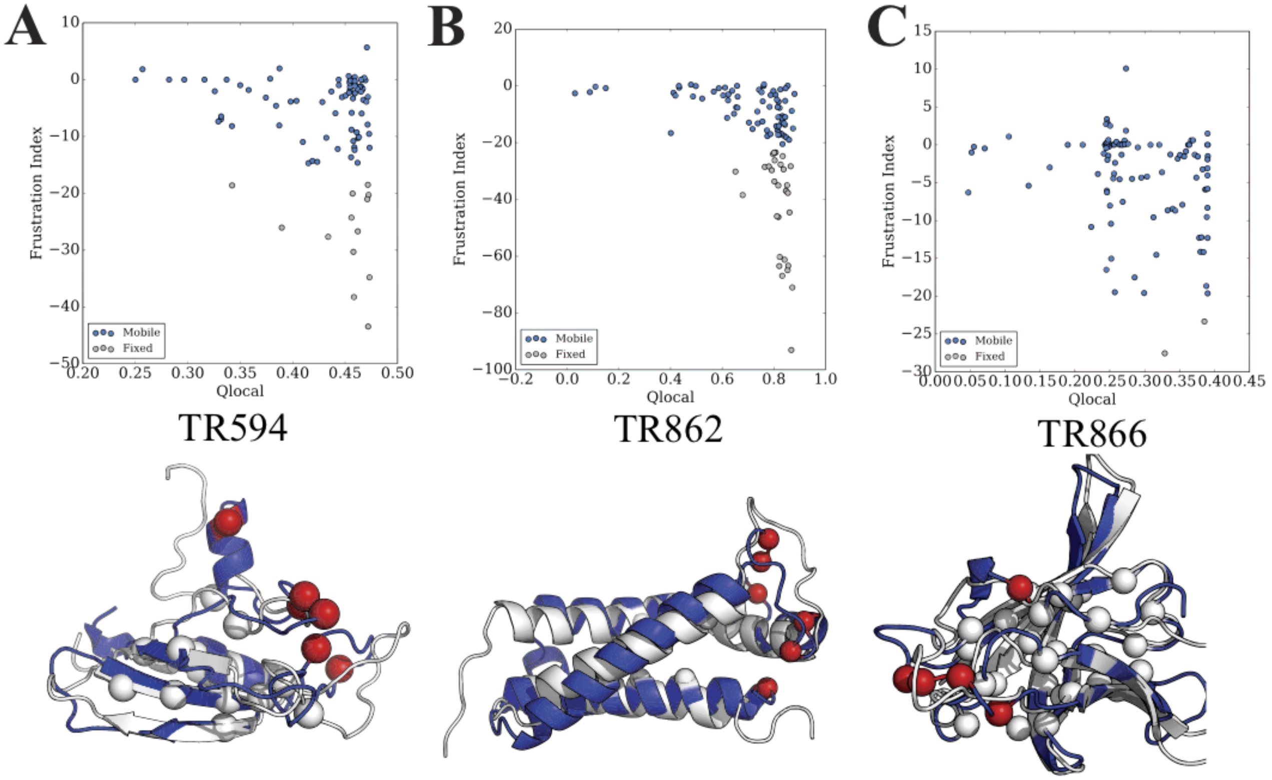


Figure S5: **Three examples from CASP12 show the relationship between the frustration pattern and local structural accuracy of the initially predicted structures** A) TR594, B) TR862, C) TR866. The upper panel shows the relationship between the local frustration pattern evaluated by frustration indices and local structural similarity by $Q_{local}$. In the lower panel, the initially predicted structure, colored by blue, was aligned to the crystal structure colored by white. The most minimally frustrated residues in the initially predicted structure which were constrained in refinement simulations are shown in white spheres. The most highly frustrated residues in the initially predicted structure are shown in red spheres.

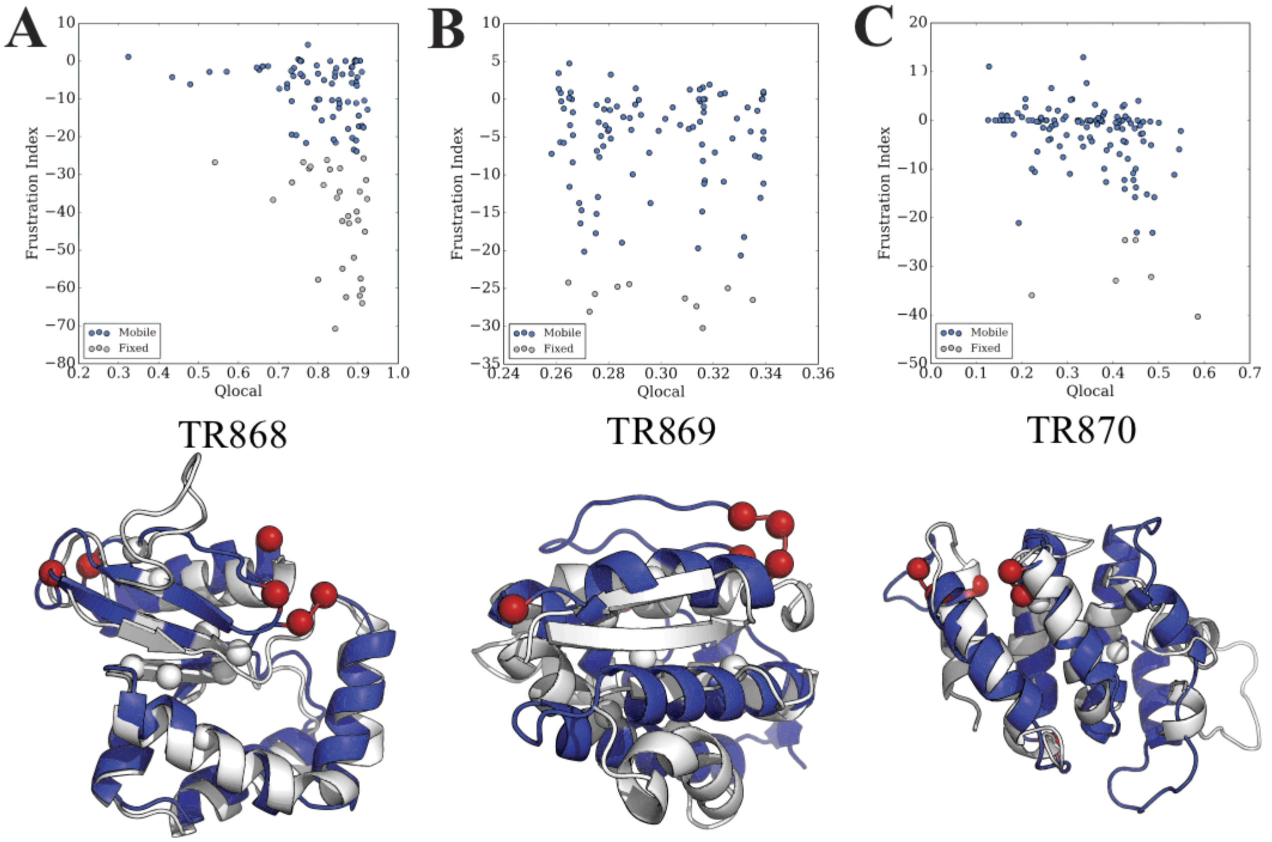


Figure S6: **Three examples from CASP12 show the relationship between the frustration pattern and local structural accuracy of the initially predicted structures** A) TR868, B) TR869, C) TR870. The upper panel shows the relationship between the local frustration pattern evaluated by frustration indices and local structural similarity by $Q_{local}$. In the lower panel, the initially predicted structure, colored by blue, was aligned to the crystal structure colored by white. The most minimally frustrated residues in the initially predicted structure which were constrained in refinement simulations are shown in white spheres. The most highly frustrated residues in the initially predicted structure are shown in red spheres.

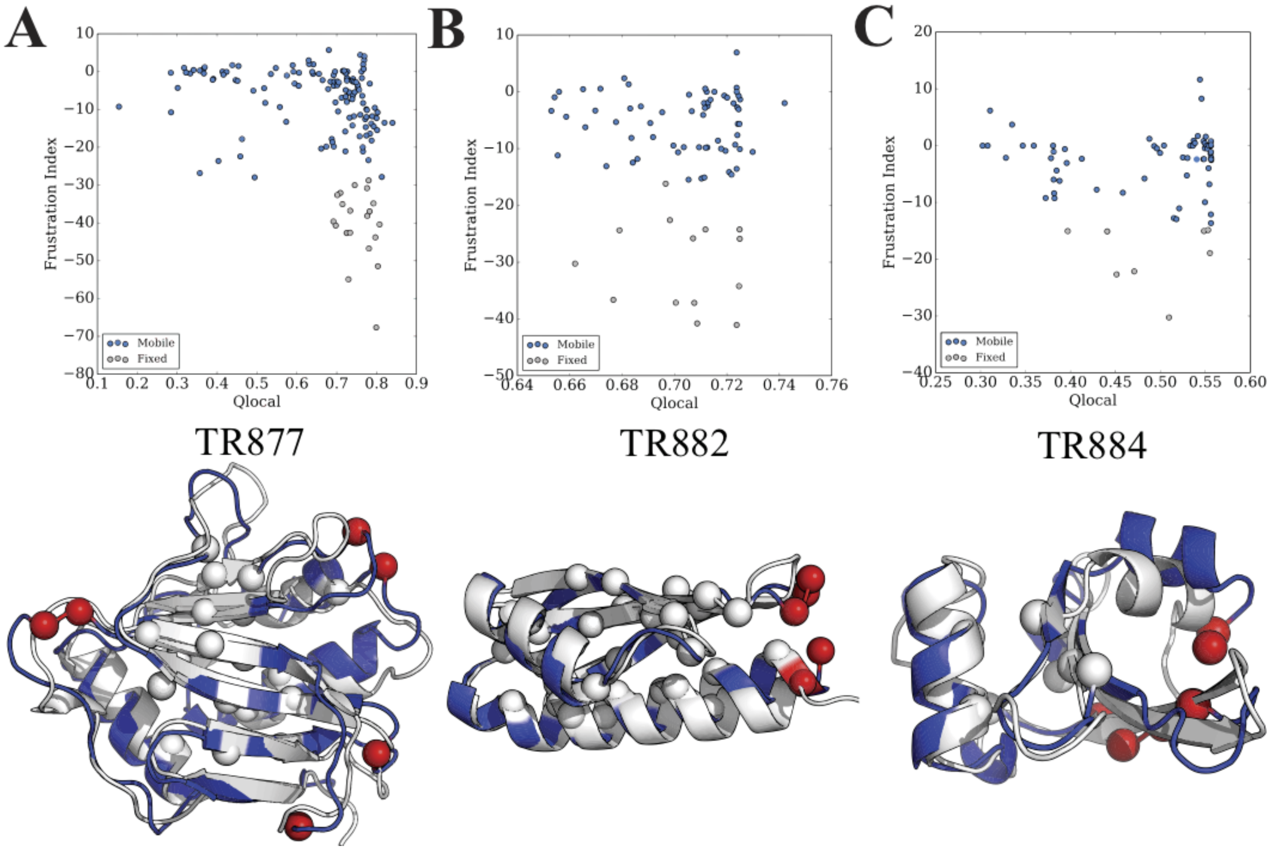


Figure S7: **Three examples from CASP12 show the relationship between the frustration pattern and local structural accuracy of the initially predicted structures** A) TR877 ,B)TR882, C) TR884. The upper panel shows the relationship between the local frustration pattern evaluated by frustration indices and local structural similarity by $Q_{local}$. In the lower panel, the initially predicted structure, colored by blue, was aligned to the crystal structure colored by white. The most minimally frustrated residues in the initially predicted structure which were constrained in refinement simulations are shown in white spheres. The most highly frustrated residues in the initially predicted structure are shown in red spheres.

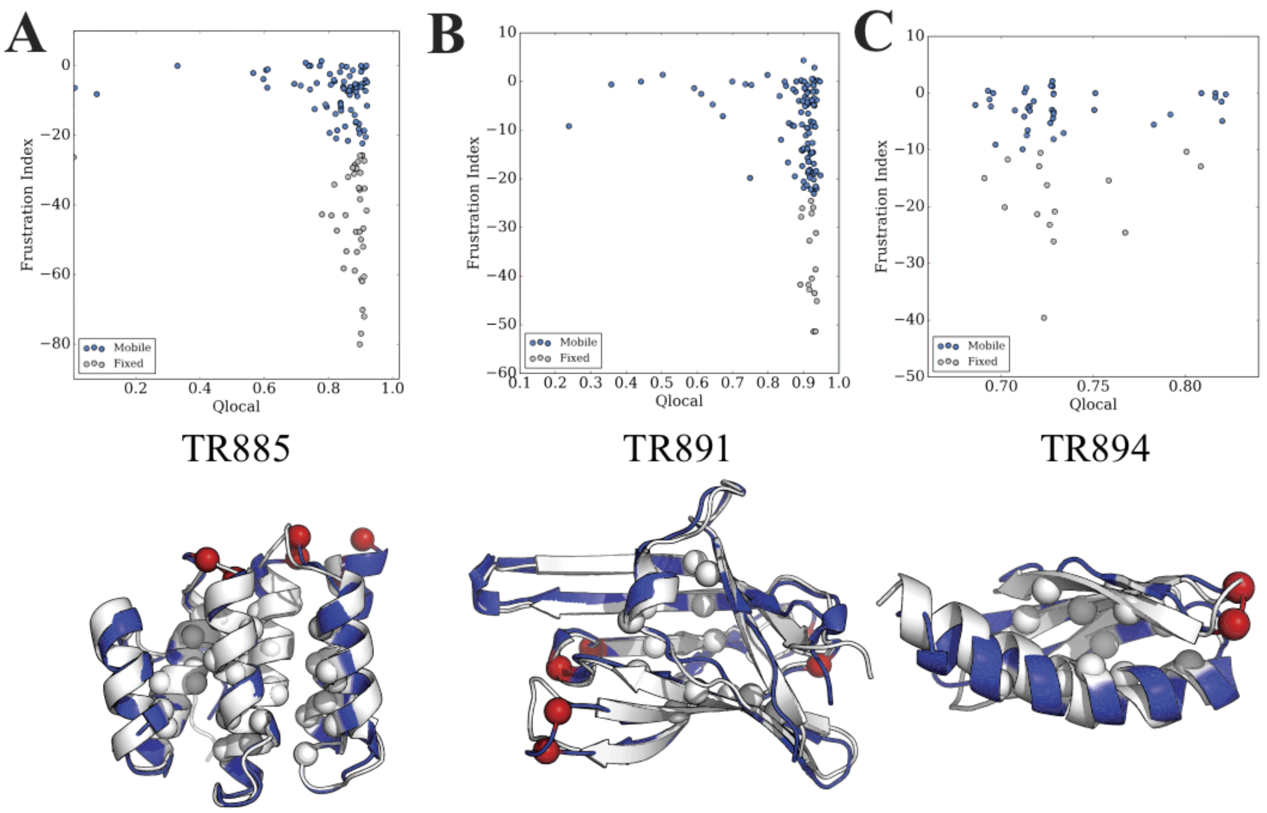


Figure S8: **Three examples from CASP12 show the relationship between the frustration pattern and local structural accuracy of the initially predicted structures** A) TR885, B) TR891, C) TR894. The upper panel shows the relationship between the local frustration pattern evaluated by frustration indices and local structural similarity by $Q_{local}$. In the lower panel, the initially predicted structure, colored by blue, was aligned to the crystal structure colored by white. The most minimally frustrated residues in the initially predicted structure which were constrained in refinement simulations are shown in white spheres. The most highly frustrated residues in the initially predicted structure are shown in red spheres.

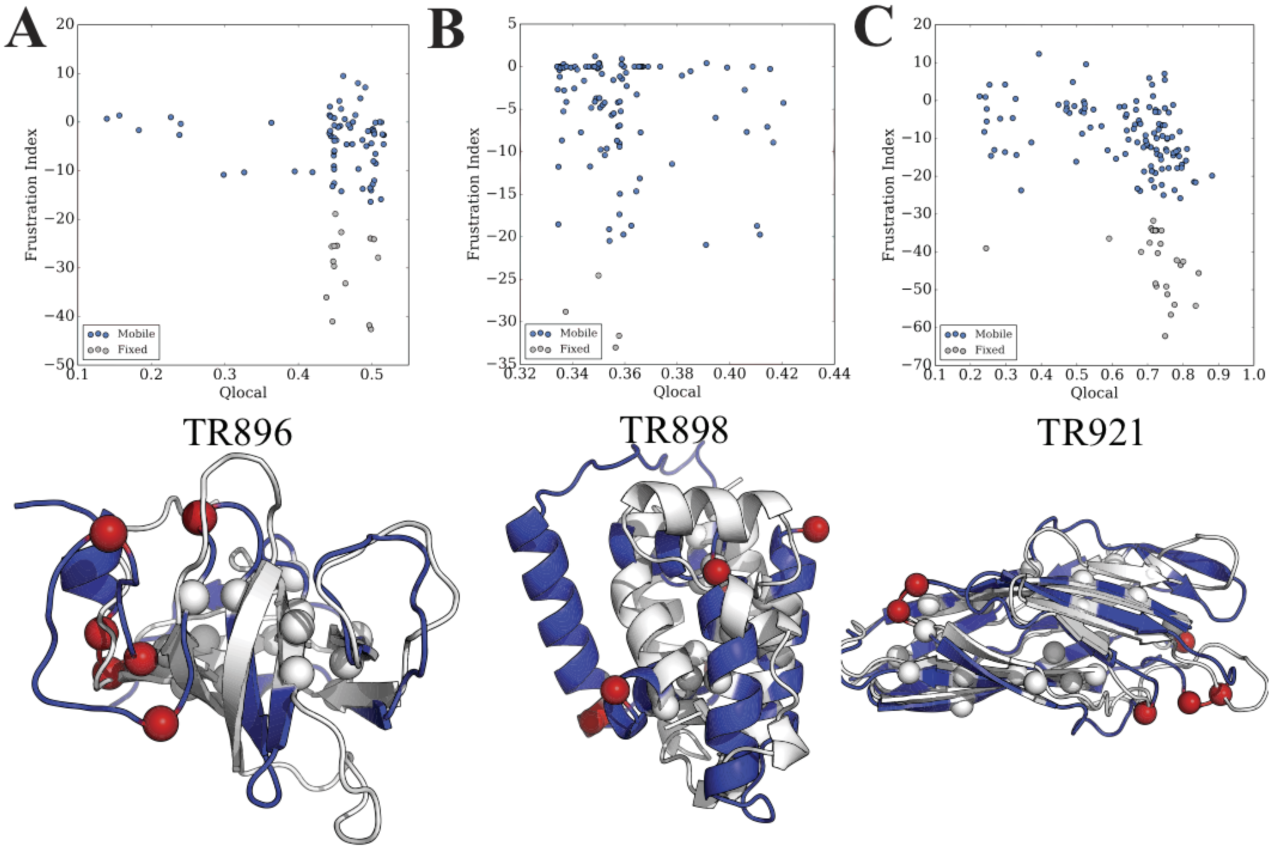


Figure S9: **Three examples from CASP12 show the relationship between the frustration pattern and local structural accuracy of the initially predicted structures** A) TR896, B) TR8,98 C)TR921. The upper panel shows the relationship between the local frustration pattern evaluated by frustration indices and local structural similarity by $Q_{local}$. In the lower panel, the initially predicted structure, colored by blue, was aligned to the crystal structure colored by white. The most minimally frustrated residues in the initially predicted structure which were constrained in refinement simulations are shown in white spheres. The most highly frustrated residues in the initially predicted structure are shown in red spheres.

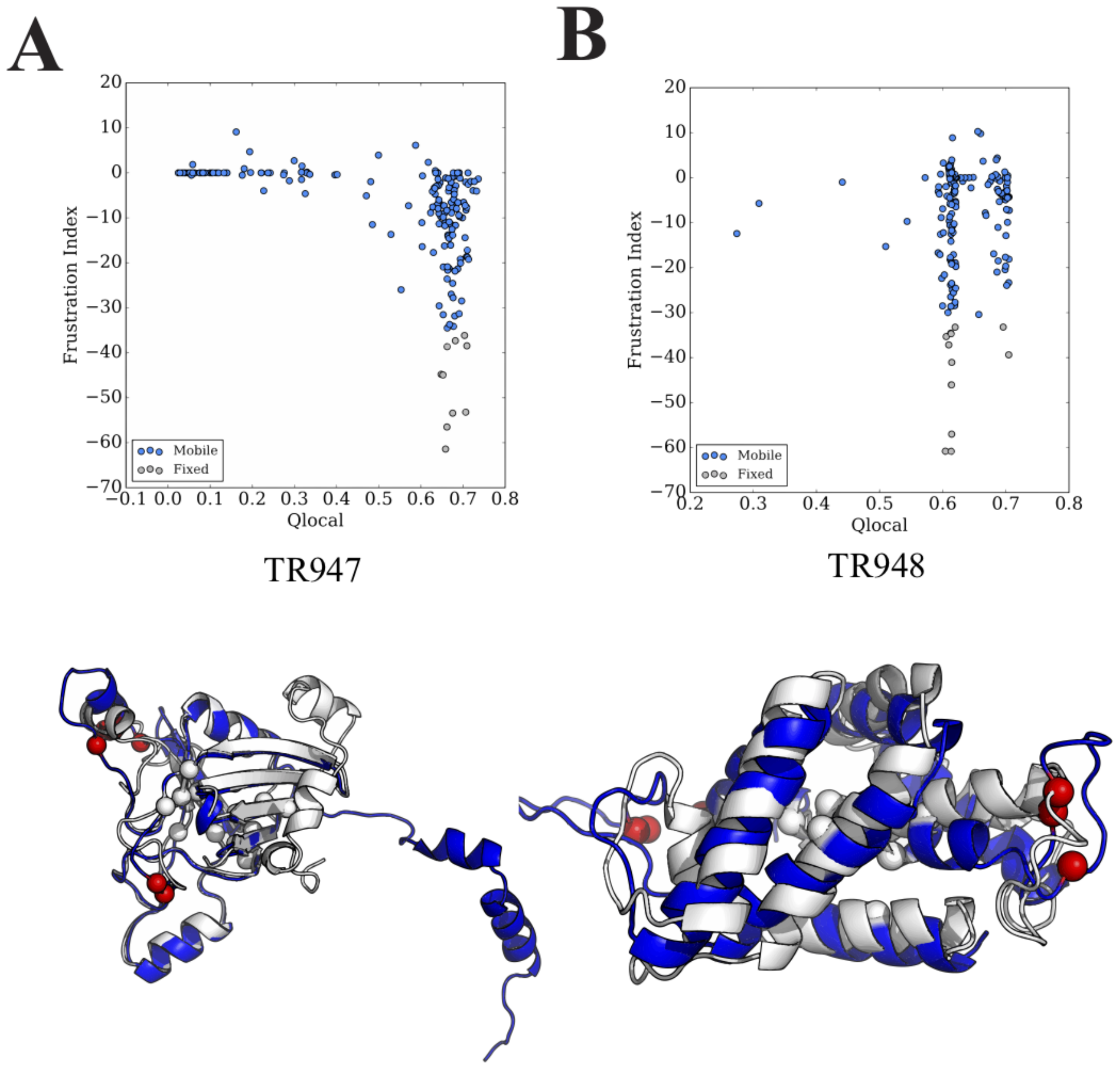


Figure S10: **Two examples from CASP12 show the relationship between the frustration pattern and local structural accuracy of the initially predicted structures** A) TR947, B) TR948. The upper panel shows the relationship between the local frustration pattern evaluated by frustration indices and local structural similarity by $Q_{local}$. In the lower panel, the initially predicted structure, colored by blue, was aligned to the crystal structure colored by white. The most minimally frustrated residues in the initially predicted structure which were constrained in refinement simulations are shown in white spheres. The most highly frustrated residues in the initially predicted structure are shown in red spheres.

#### S3 The change of sidechain accuracy during simulation

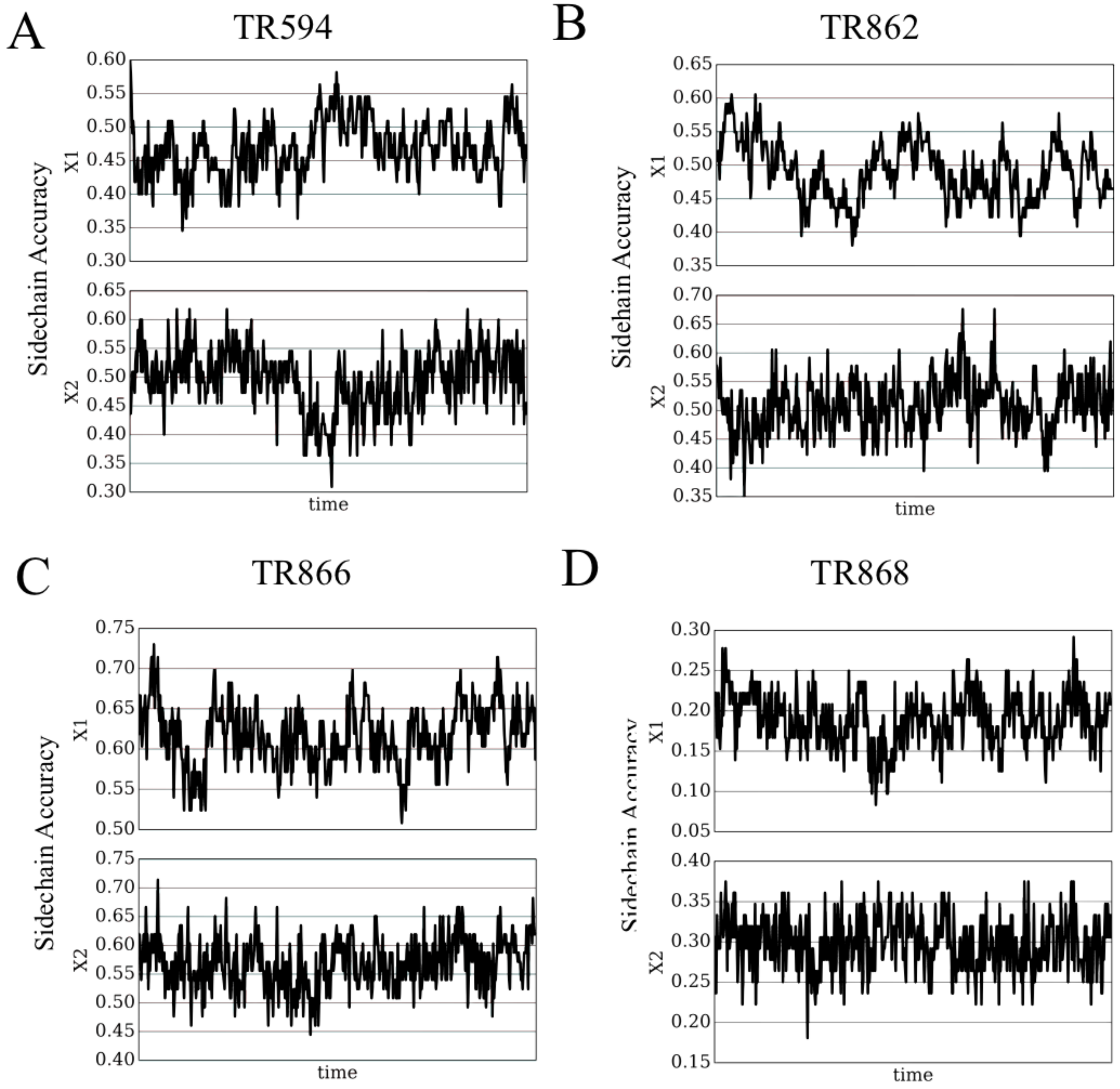


Figure S11: **The change of sidechain accuracy during refinement simulation** A) TR594, B) TR862, C) TR866, D) TR868. X1 X2 are shown in these figures

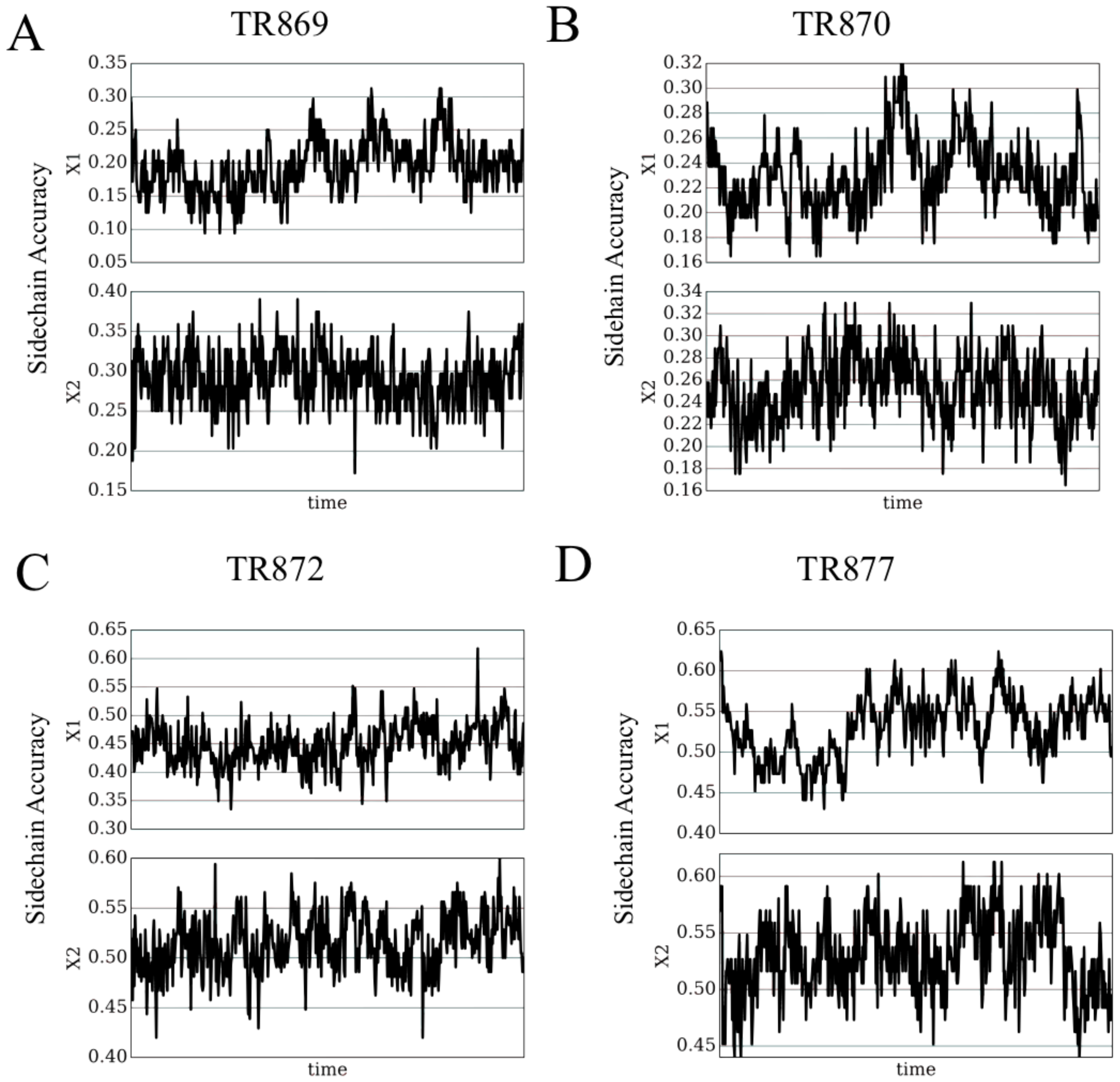


Figure S12: **The change of sidechain accuracy during refinement simulation** A) TR869, B) TR870, C) TR872, D) TR877. X1 X2 are shown in these figures

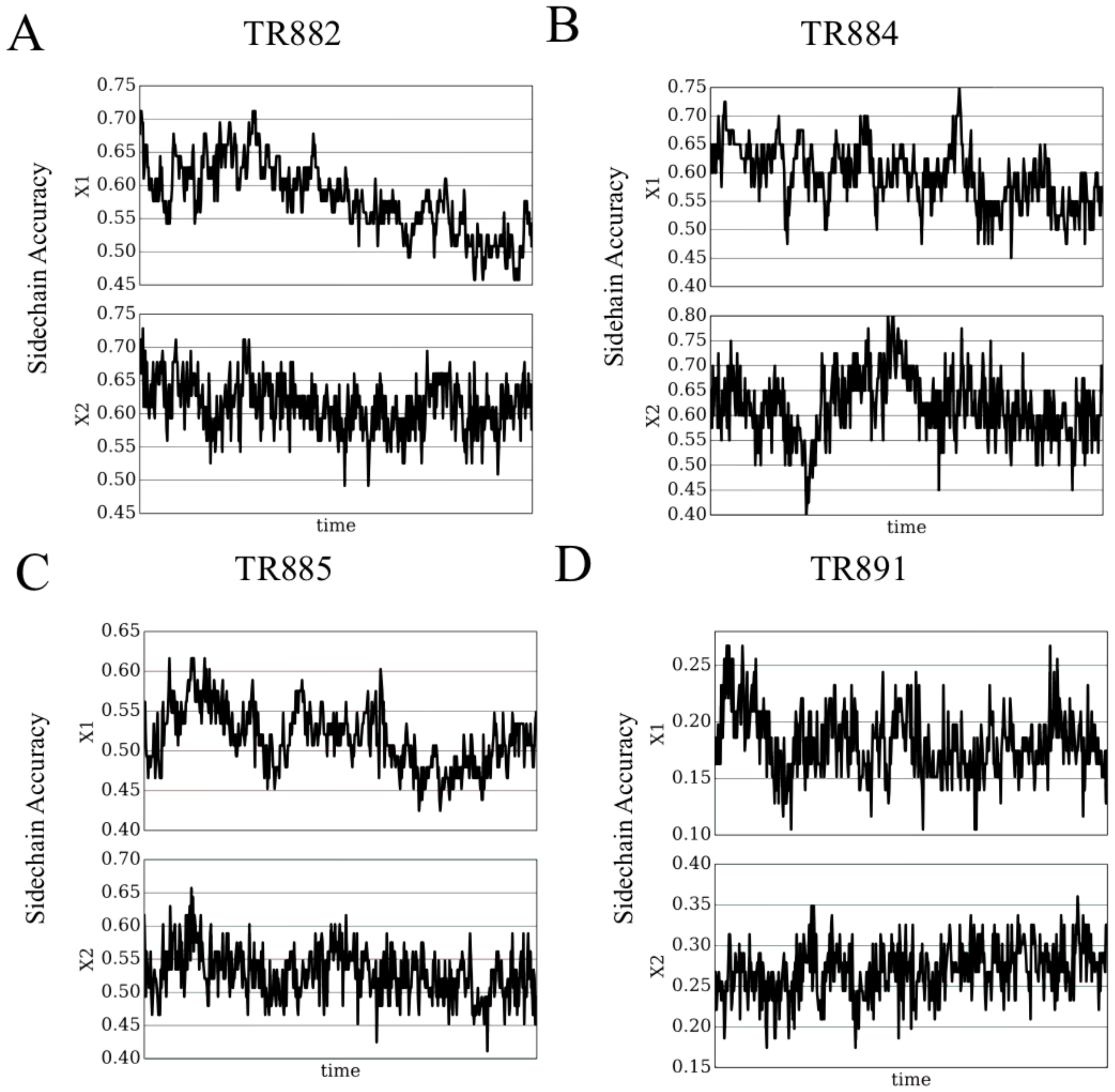


Figure S13: **The change of sidechain Accuracy during refinement simulation** A) TR882, B) TR884, C) TR885, D) TR891. X1 X2 are shown in these figures

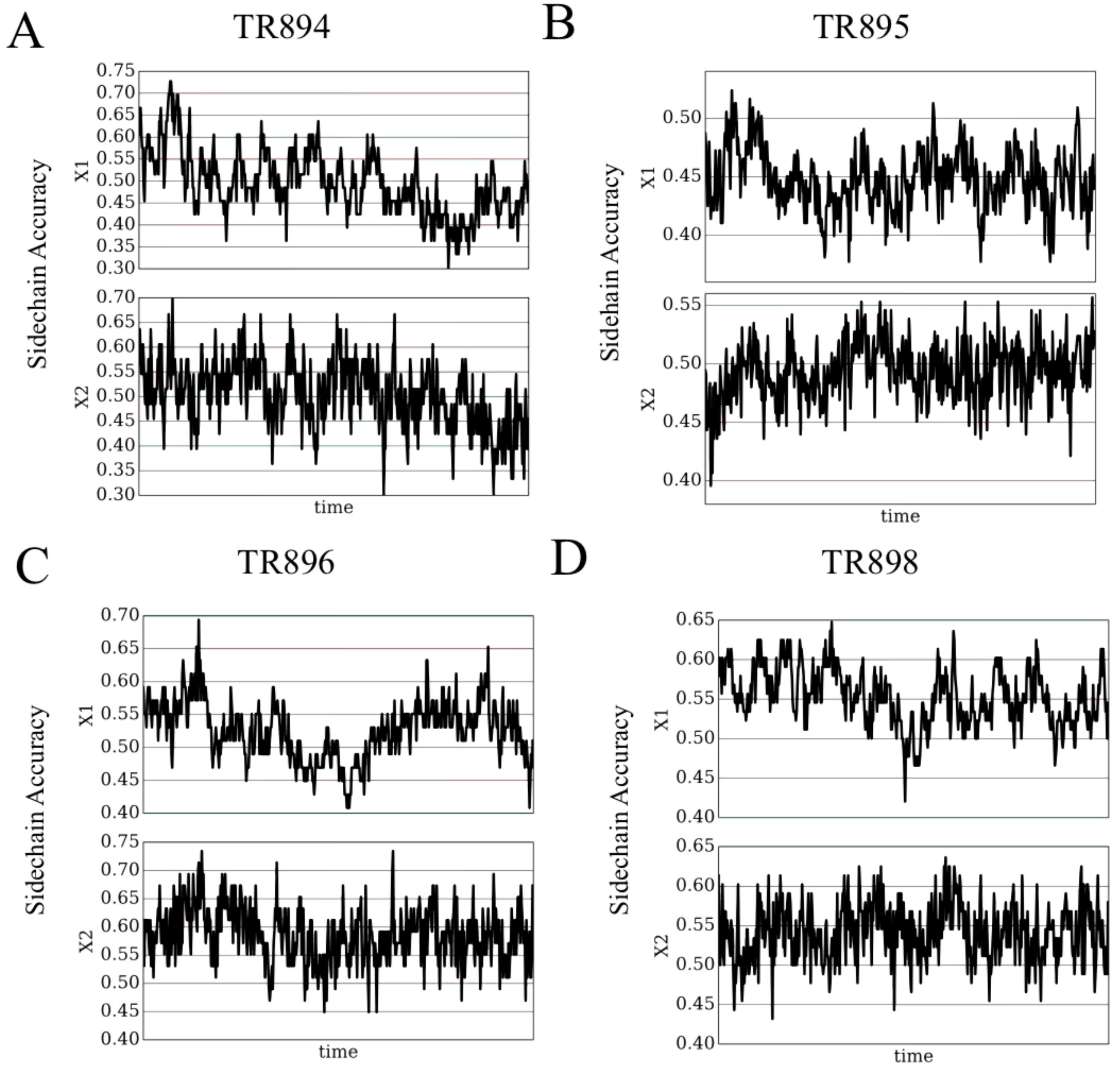


Figure S14: **The change of sidechain accuracy during refinement simulation** A) TR894, B) TR895, C) TR896, D) TR898. X1 X2 are shown in these figures

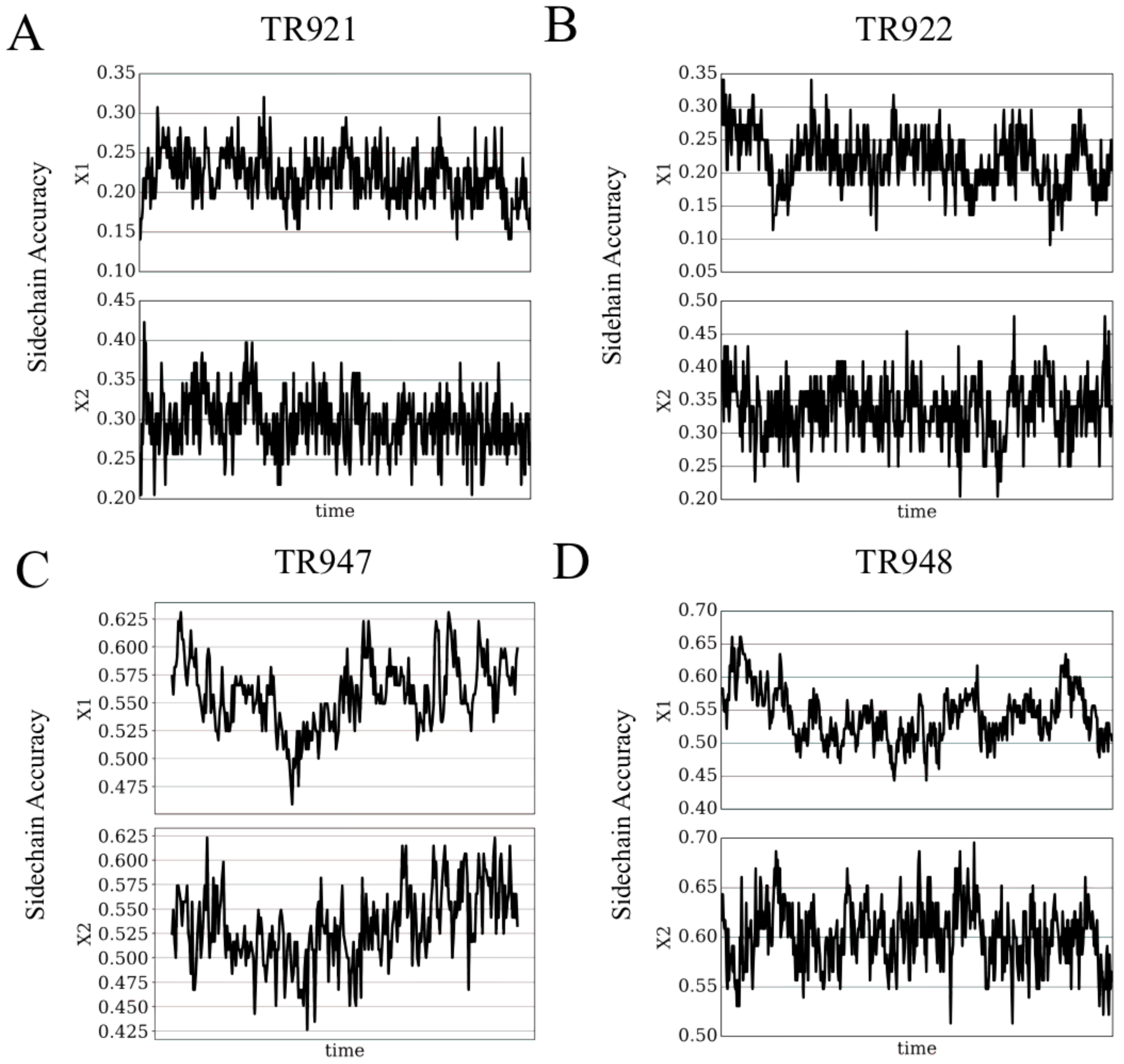


Figure S15: **The change of sidechain accuracy during refinement simulation** A) TR921, B) TR922, C) TR947, D) TR948. X1 X2 are shown in these figures

#### S4 Refinements guided by atomic packing frustration guides the structures towards its experimentally determined native states

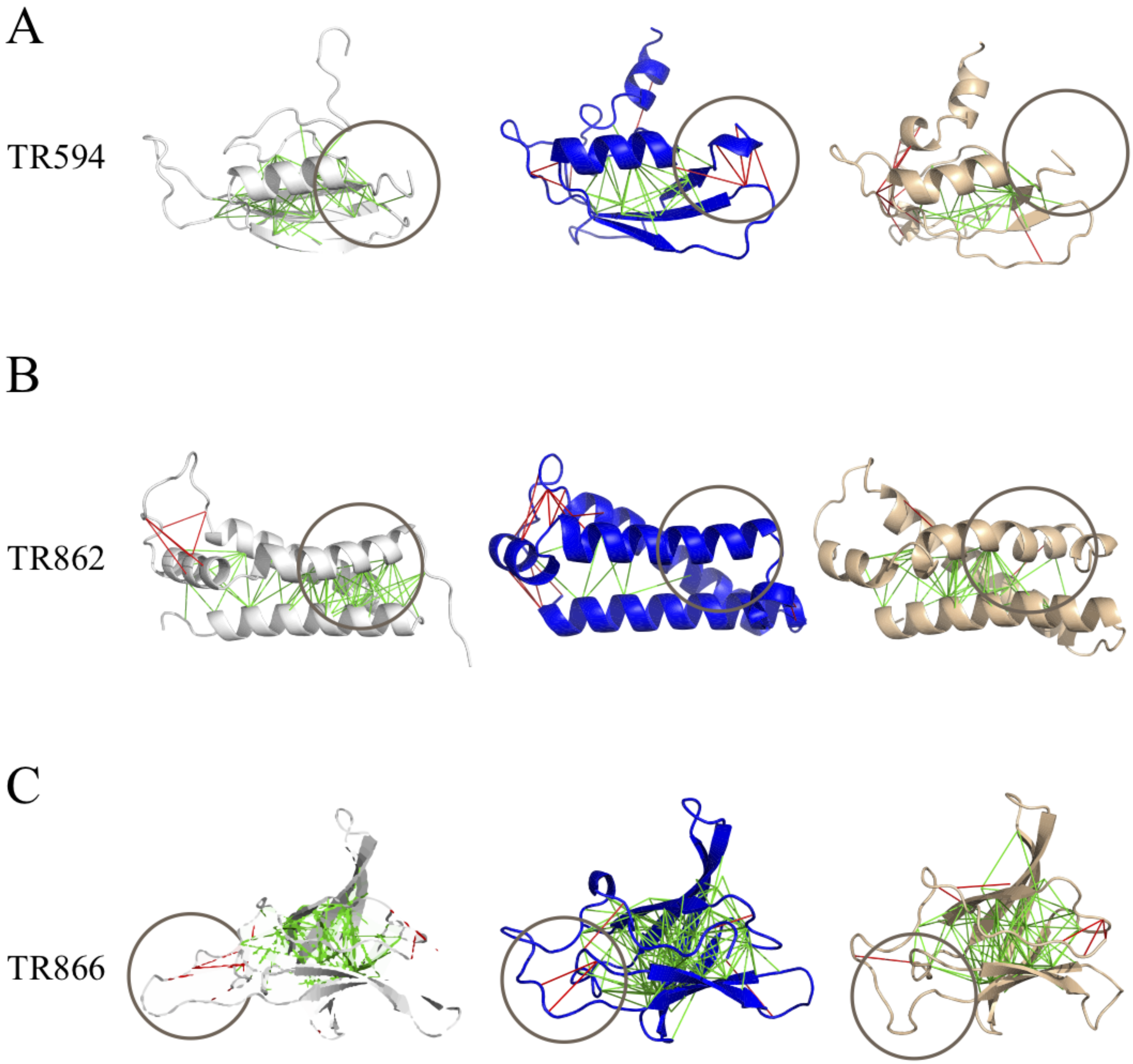


Figure S16: **Differences of the frustration patterns between the crystal structures, the initially predicted structures and the refined structure obtained by blind selection** A) TR594, B) TR862, C) TR866. The left panel shows the frustration pattern of the crystal structures, the middle panel shows the frustration pattern of the initially predicted structures, and the right panel shows the frustration pattern of the selected structures from refinement. The green lines indicate minimally frustrated interactions and the red lines indicate highly frustrated interactions. The gray circle highlights the regions where the RMSD values and the amount of highly frustrated interactions decrease from those of the initially predicted structures, which suggests the effectiveness of atomic packing frustration guided refinement.

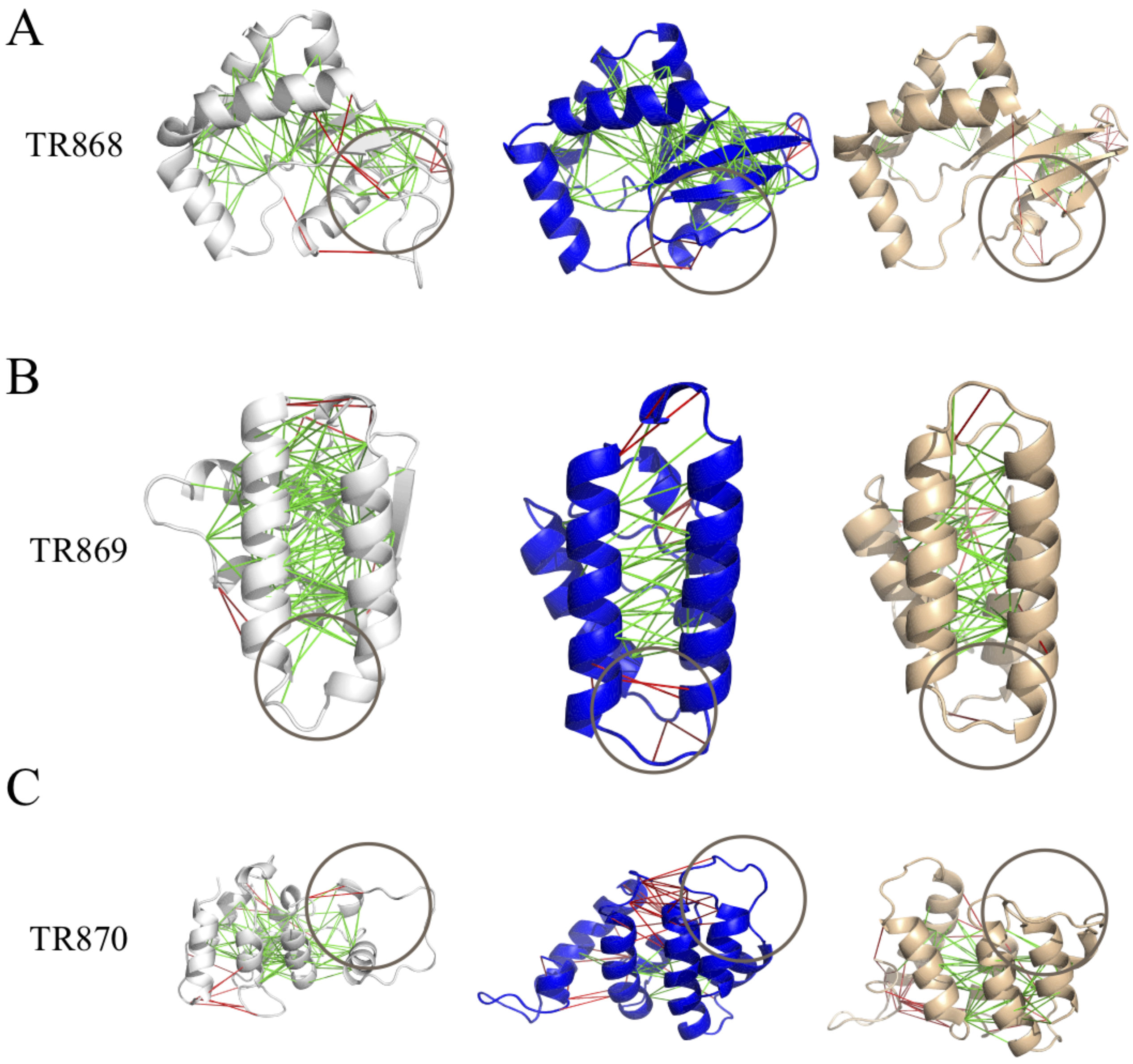


Figure S17: **Differences of the frustration patterns between the crystal structures, the initially predicted structures and the refined structure obtained by blind selection** A) TR868, B) TR869, C) TR870. The left panel shows the frustration pattern of the crystal structures, the middle panel shows the frustration pattern of the initially predicted structures, and the right panel shows the frustration pattern of the selected structures from refinement. The green lines indicate minimally frustrated interactions and the red lines indicate highly frustrated interactions. The gray circle highlights the regions where the RMSD values and the amount of highly frustrated interactions decrease from those of the initially predicted structures, which suggests the effectiveness of atomic packing frustration guided refinement.

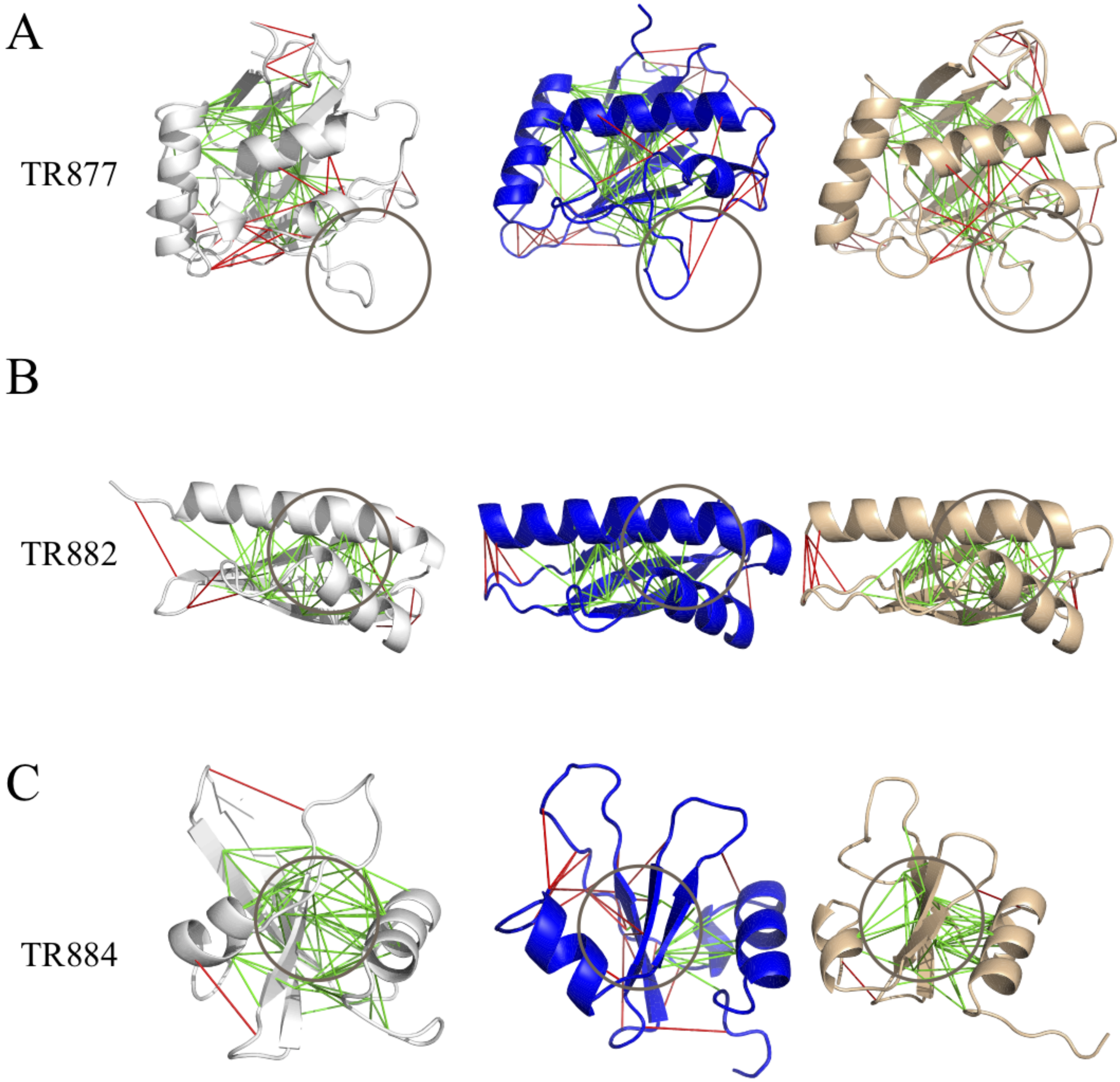


Figure S18: **Differences of the frustration patterns between the crystal structures, the initially predicted structures and the refined structure obtained by blind selection** A) TR877, B) TR882, C) TR884. The left panel shows the frustration pattern of the crystal structures, the middle panel shows the frustration pattern of the initially predicted structures, and the right panel shows the frustration pattern of the selected structures from refinement. The green lines indicate minimally frustrated interactions and the red lines indicate highly frustrated interactions. The gray circle highlights the regions where the RMSD values and the amount of highly frustrated interactions decrease from those of the initially predicted structures, which suggests the effectiveness of atomic packing frustration guided refinement.

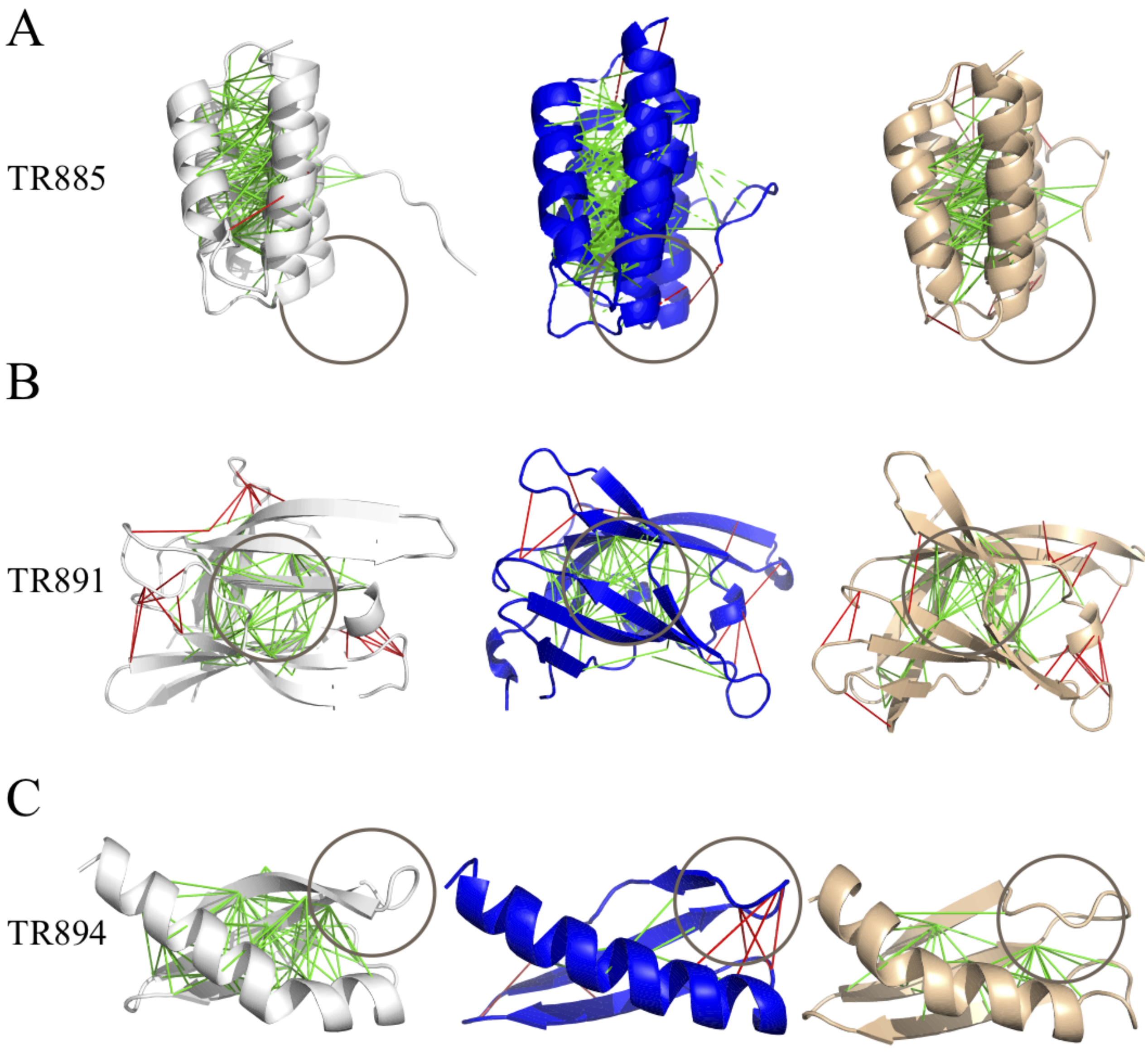


Figure S19: **Differences of the frustration patterns between the crystal structures, the initially predicted structures and the refined structure obtained by blind selection** A) TR885, B) TR891, C) TR894. The left panel shows the frustration pattern of the crystal structures, the middle panel shows the frustration pattern of the initially predicted structures, and the right panel shows the frustration pattern of the selected structures from refinement. The green lines indicate minimally frustrated interactions and the red lines indicate highly frustrated interactions. The gray circle highlights the regions where the RMSD values and the amount of highly frustrated interactions decrease from those of the initially predicted structures, which suggests the effectiveness of atomic packing frustration guided refinement.

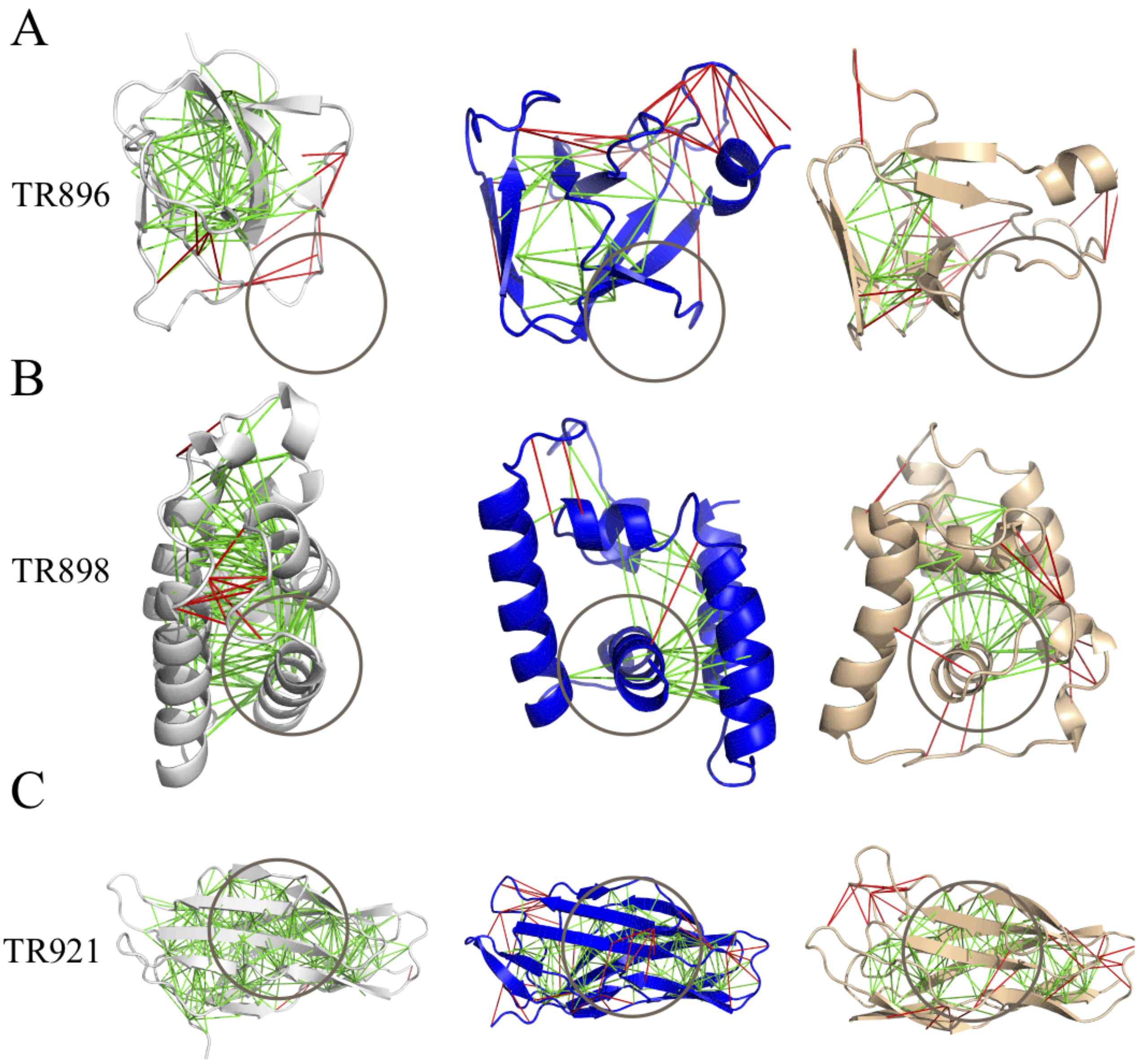


Figure S20: **Differences of the frustration patterns between the crystal structures, the initially predicted structures and the refined structure obtained by blind selection** A) TR896, B) TR898, C) TR921. The left panel shows the frustration pattern of the crystal structures, the middle panel shows the frustration pattern of the initially predicted structures, and the right panel shows the frustration pattern of the selected structures from refinement. The green lines indicate minimally frustrated interactions and the red lines indicate highly frustrated interactions. The gray circle highlights the regions where the RMSD values and the amount of highly frustrated interactions decrease from those of the initially predicted structures, which suggests the effectiveness of atomic packing frustration guided refinement.

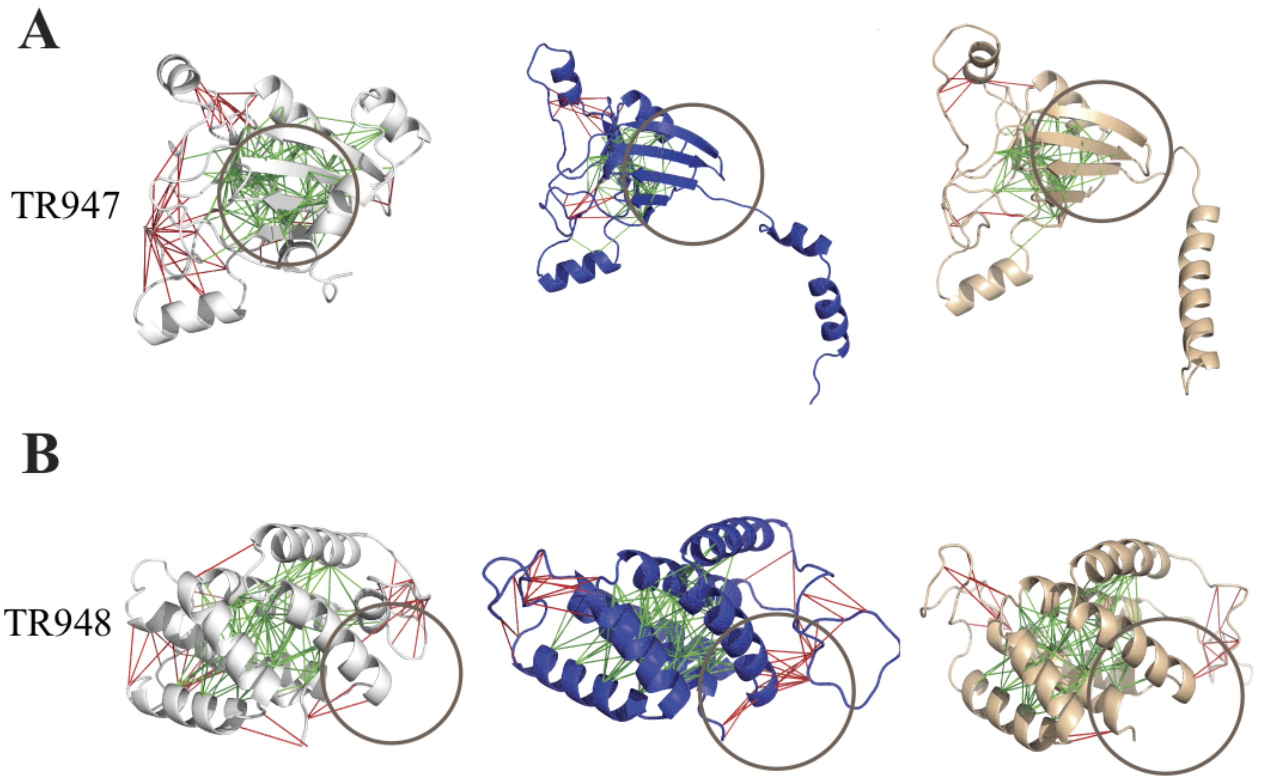


Figure S21: **Differences of the frustration patterns between the crystal structures, the initially predicted structures and the refined structure obtained by blind selection** A) TR947, B) TR948. The left panel shows the frustration pattern of the crystal structures, the middle panel shows the frustration pattern of the initially predicted structures, and the right panel shows the frustration pattern of the selected structures from refinement. The green lines indicate minimally frustrated interactions and the red lines indicate highly frustrated interactions. The gray circle highlights the regions where the RMSD values and the amount of highly frustrated interactions decrease from those of the initially predicted structures, which suggests the effectiveness of atomic packing frustration guided refinement.

#### S5 Comparing performance of atomic packing frustration guided refinement with other refinement methods by GDT-TS score

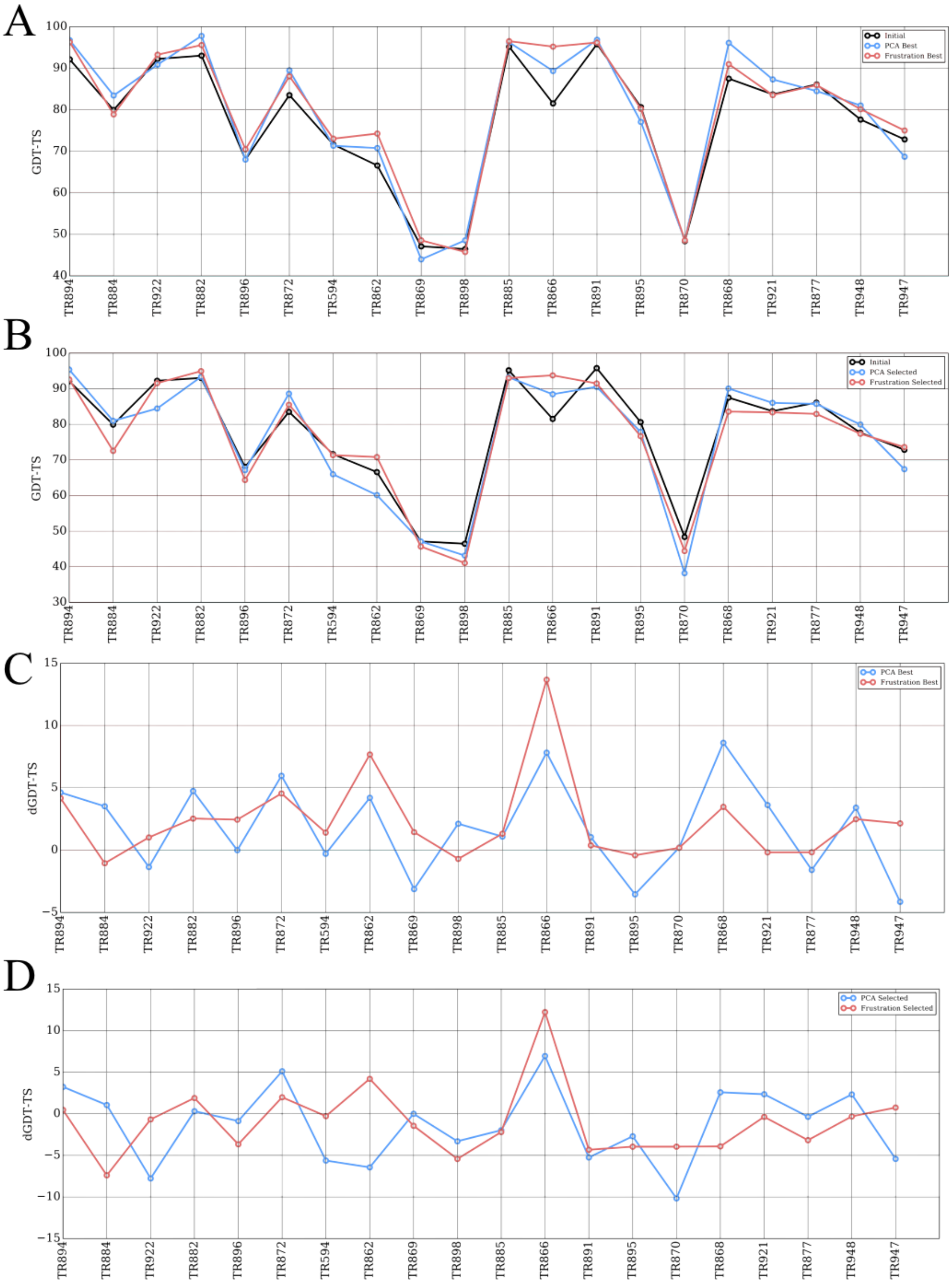


Figure S22: **A summary of refinement results. A)The GDT-TS score of initially predicted and refined structures with the highest GDT-TS score** Initial: Black, PC-guided refinement: Blue, Atomic Packing Frustration Refinement: Green. B)The GDT-TS score of initially predicted structure and selected structures from different refinement strategy: Initial: Black,PC-guided refinement: Blue, Atomic Packing Frustration Refinement: Green.B)The ΔGDT-TS score between initially predicted structures and refined structures with the highest GDT-TS score:PC-guided refinement: Blue, Atomic Packing Frustration Refinement: Green. B)The ΔGDT-TS score between initially predicted structure and selected structures from different refinement strategy: PC-guided refinement: Blue, Atomic Packing Frustration Refinement: Green.

#### The Atomic Packing Frustration Guided Refinement Drive the Structures to the Native State

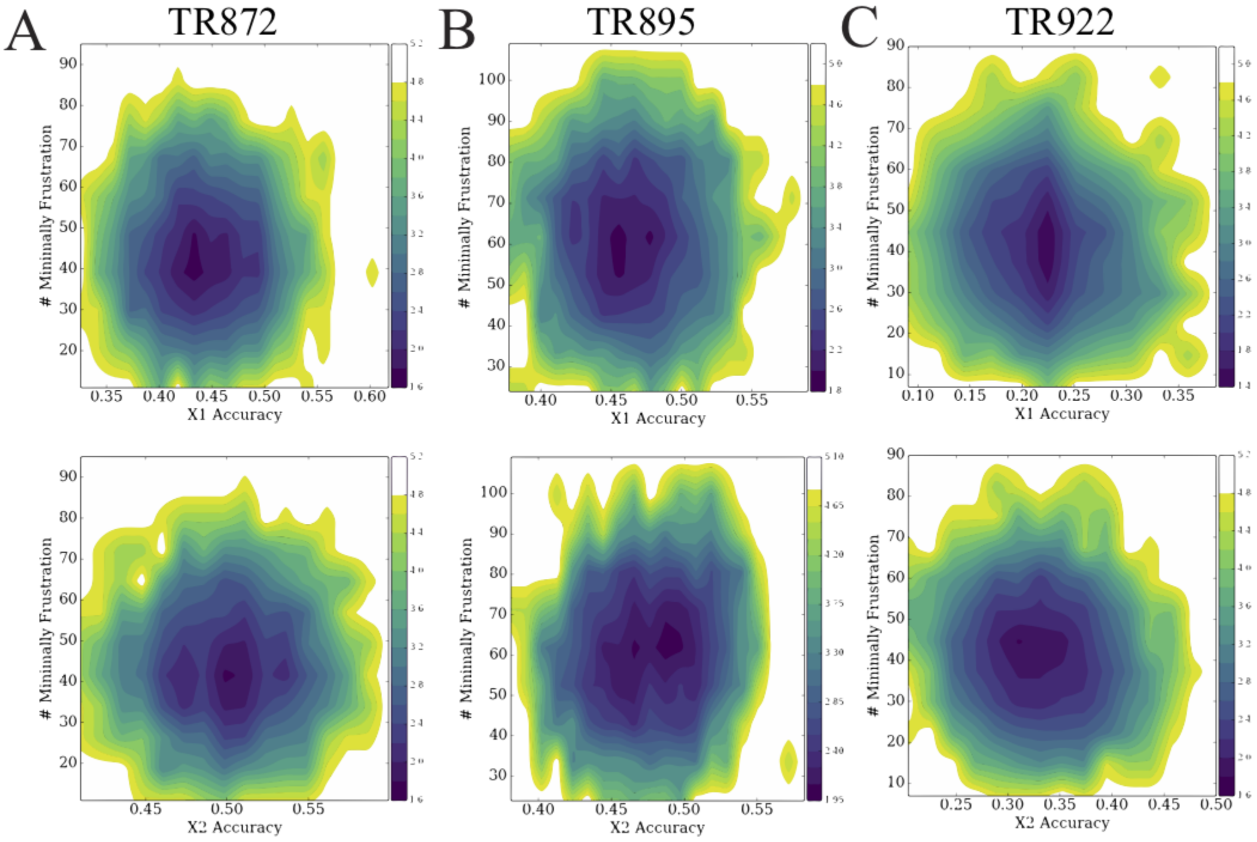


Figure S23: **The free energy profiles projected to the number of minimally frustrated interaction and X1 or X2 Accuracy** A)T0872,B)T0895, C)T0922.

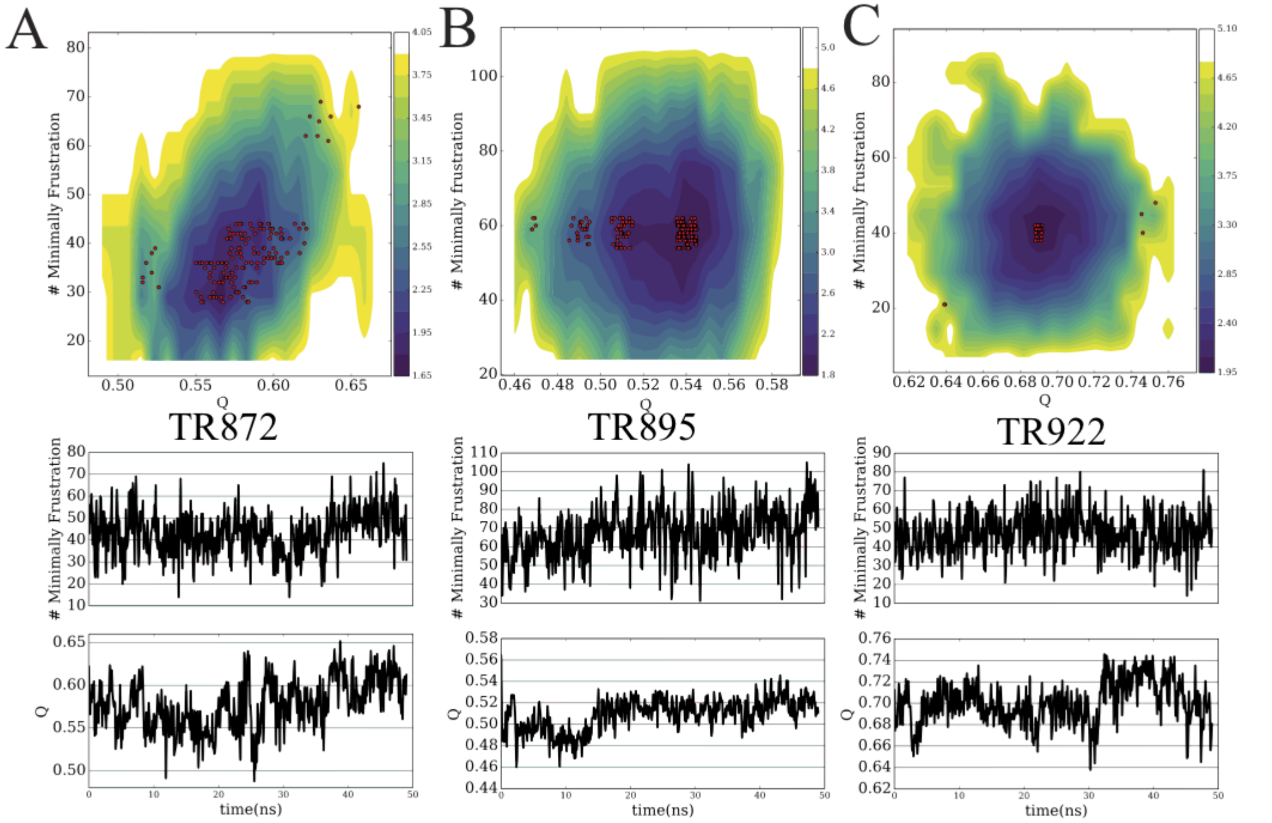


Figure S24: **The summary of refinement processes** A)T0872, B)T0895, C) T0922. The upper panel shows free energy profiles using the number of minimally frustrated interactions and q value as two dimensions. The red dots are some structures which suggest the trajectory of refine process. The lower panel gives the change of the number of minimally frustrated interaction and q in the simulations.

### S6 The free-energy landscapes of TR872 at temperature 300K

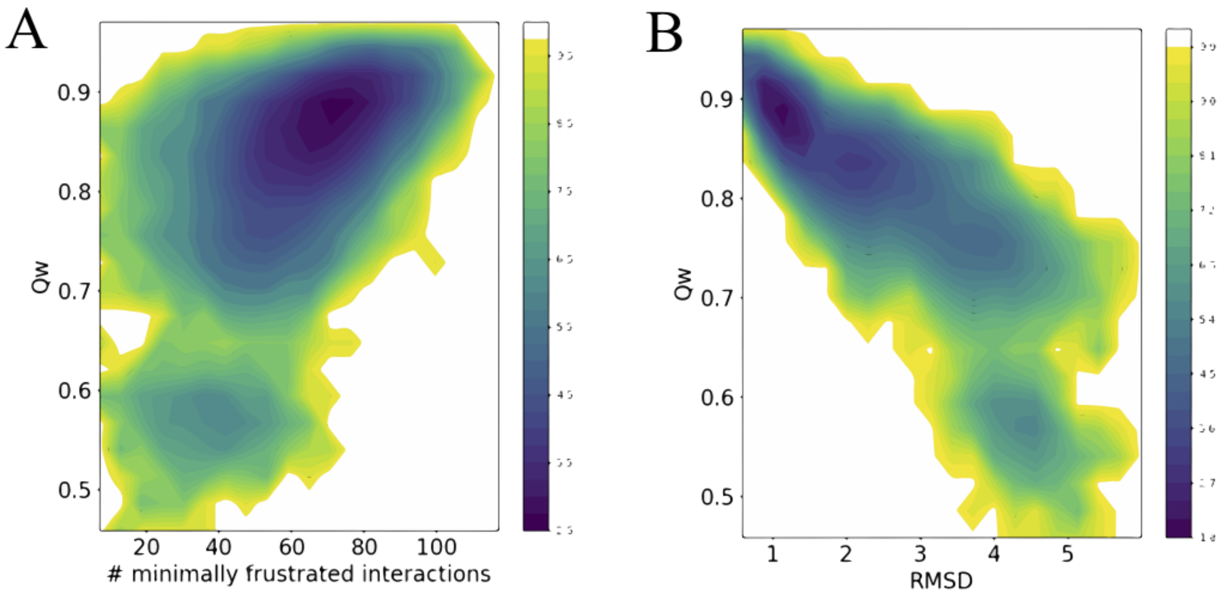


Figure S25: **The free-energy profiles of TR872 at Temperature 300K** A) The 2D free-energy surface is plotted using the number of minimally frustrated interactions and the Qw value. B)The 2D free-energy profile is plotted using RMSD value and the Qw value.
